# Continental-scale genomic surveillance of *Plasmodium falciparum* malaria with rapid nanopore sequencing

**DOI:** 10.1101/2025.07.23.666274

**Authors:** Mulenga Mwenda, Karolina Mosler, Bernd Bohmeier, Miriam Chomba, Welmoed Van Loon, Brenda Mambwe, Amy Gaye, Adedolapo Olorunfemi, Salma Suliman, Nassandba Julien Yanogo, Djiby Sow, Bassirou Ngom, Oumou Aïcha Zeïna Zoure, Yssimini Nadège Guillène Tibiri, Fiyinfoluwa Ojeniyi, Arsène Zongo, Etilé A. Anoh, Vincent Achi, Carol Chiyesu, Sheila Otieno, Bixa Ogola, Moussa Niangaly, Manuela Carrasquilla, Dagaga Kenea Goboto, Torsten Feldt, Tafese Beyene Tufa, Rafael Oliveira, Emma Schallenberg, Yuhana Sogoba, Christina Ntalla, Oumarou Ouedraogo, Kephas Otieno, Oluyinka Opaleye, Adekunle Olowe, Marley Gibbons, Chris Drakeley, Grit Schubert, Frank P. Mockenhaupt, Silvia Portugal, Awa B. Deme, Issiaka Soulama, Daouda Ndiaye, Olusola Ojurongbe, Simon Kariuki, Ya Ping Shi, Jonathan S. Schultz, Moonga Hawela, Daniel J. Bridges, Jason A. Hendry

## Abstract

In sub-Saharan Africa, continental-scale genomic surveillance of *Plasmodium falciparum* malaria is needed to track the spread of antimalarial drug resistance and diagnostic test evasion, as well as to monitor parasite evolutionary responses to vaccine rollout. Yet implementation of malaria genomic surveillance at a continental-scale is hindered by resource constraints, the vastness of the continent, and the lack of sequencing protocols suitable for most local laboratories. To address this, we developed an approach to enable a decentralized scale-up of *P. falciparum* genomic surveillance and established it in six African countries in one year, locally sequencing 1,065 samples. The approach includes a rapid (∼ 5 hours) and cost-efficient (<$25 USD/sample) nanopore sequencing protocol that provides surveillance of drug resistance-associated genes, *hrp2/3* deletions, the vaccine target *csp*, and the polymorphic gene *ama1*. We coupled this to a bioinformatics dashboard that runs offline on a laptop and displays mapping and variant calling results in real-time. We demonstrate robust sequencing coverage across parasitemia levels and laboratories, accurate identification of antimalarial resistance markers and *hrp2/3* deletions; and, with a novel variant caller, sensitive detection of mutations carried by minor clones. Our approach will accelerate genomic surveillance of *P. falciparum* malaria across sub-Saharan Africa at a time of urgent need.

## Introduction

The endeavor to eliminate *Plasmodium (P*.*) falciparum* malaria from sub-Saharan Africa – where it causes over half a million deaths annually – relies heavily on accurate diagnosis with rapid diagnostic tests (RDTs) and effective treatment with artemisinin-based combination therapies (ACTs)^1^. *P. falciparum*, however, is evolving to undermine both of these tools. For RDTs, *P. falciparum* strains that have deletions of the *hrp2* and *hrp3* genes evade diagnosis by producing false-negative test results^2–6^. For ACTs, mutations in the *kelch13* gene^7^ can cause delayed parasite clearance^8^ and, if combined with partner drug resistance, treatment failure^9^. Since at least 2016, these *kelch13* mutations have been spreading in East Africa^10–13^, where they have begun to co-circulate with *hrp2/3* deletions^14^. They are now being described in Southern^15^ and Central Africa^16,17^. Public health institutions are urgently exploring a variety of responses, such as switching to alternative RDTs when the *hrp2/3* deletion frequency surpasses a threshold regionally^18^; or deploying multiple first-line therapies (MFT) to mitigate the spread of *kelch13* mutations^19,20^. However, essential to coordinating these responses are high-quality, granular, and timely malaria surveillance data, which are lacking in many parts of Africa.

The need to increase malaria surveillance in Africa could be met using genomics. Genomic methods like amplicon sequencing^21–23^ and molecular inversion probes (MIPs)^24,25^ can interrogate tens to thousands of *P. falciparum* genes in parallel, enabling reporting on many control-relevant genetic markers at once. This makes them more efficient than conventional molecular methods, which typically only report on one gene or mutation. Moreover, many genomic methods require just a single dried-blood spot (DBS) as the sample, making them less invasive and labour intensive than standard surveillance approaches like therapeutic efficacy studies. In principle, these genomics approaches could affordably and routinely generate high-quality malaria surveillance data. But scaling *P. falciparum* genomic surveillance to meet the growing need across sub-Saharan Africa is an immense challenge. First, the region is vast and diverse, spanning over 24 million km^2^ and encompassing 49 countries. Perhaps 50,000 to 100,000 samples would need to be sequenced annually to have adequate statistical power to detect emerging mutations across the entire region. Second, access to Next-Generation Sequencing (NGS) platforms is limited in many areas. Where NGS platforms do exist, maintenance and the procurement of reagents is invariably unreliable and expensive. Similarly, the computational infrastructure (and/or internet connectivity) required to process large volumes of genomic data is frequently lacking. As a consequence, most *P. falciparum* genomic data have been generated by shipping samples out of Africa, a slow process that also impedes local capacity development and ownership. More recently, centralised sequencing facilities located within Africa are beginning to generate malaria genomic data domestically and regionally, but questions over capacity and country ownership persist.

The possibility of scaling *P. falciparum* genomic surveillance across sub-Saharan Africa in a decentralised way has been relatively neglected. The decentralised approach would leverage the portable and low-cost MinION device from Oxford Nanopore Technologies (ONT)^26^ to conduct genomic surveillance from a network of laboratories that span the continent. This would circumvent the need for large capital investments associated with centralised facilities and Illumina platforms, and instead engage more of the existing funding, infrastructure and human potential latent in the thousands of conventional laboratories across Africa. These laboratories could serve their local geography by interacting closely with nearby clinics and public health officials; while simultaneously coordinating activities and sharing data to improve national and continental understanding. Beneficially, a network of decentralised sequencing laboratories would be more robust to disruption and could grow autonomously and exponentially. Until now, the major drawback for this paradigm has been that nanopore sequencing protocols for *P. falciparum* genomic surveillance, although increasing in number^27–30^, are less mature and comprehensive than Illumina-based ones^21–23,31^. For example, state-of-the-art Illumina-based amplicon sequencing protocols^23^ concurrently provide information on antimalarial drug resistance, *hrp2/3* deletions, the vaccine target *csp* and multiple genetic diversity markers. In contrast, existing nanopore sequencing protocols typically provide information on only one^27,28^ or two^29,30^ of these outputs. An exception is the NOMADS16 protocol^32^ but its focus on generating long-read data (3–4 kbp) results in it being substantially less sensitive than short-read approaches.

Here, we have filled this gap by developing a novel nanopore sequencing protocol for *P. falciparum* malaria that is rapid, cost-effective, sensitive, and comprehensive — providing information on a panel of genes associated with antimalarial drug resistance (*crt, dhfr, dhps, kelch13, mdr1*), as well as *hrp2/3* deletions, the vaccine target *csp*, and the highly polymorphic gene *ama1*. To enable on-site analysis, we developed a real-time bioinformatics pipeline and dashboard that can run on the same laptop used for sequencing. Using these tools, we locally sequenced and analysed 1,065 *P. falciparum* DBS samples from six laboratories spanning sub-Saharan Africa in one year. In addition, we developed a novel variant caller to enable sensitive detection of minor clones in polyclonal *P. falciparum* infections, and validate critical aspects of protocol performance across mock and field samples.

## Results

### A protocol to rapidly scale-up decentralised malaria genomic surveillance

We developed an optimal nanopore sequencing protocol for scaling decentralised *P. falciparum* genomic surveillance across sub-Saharan Africa (Fig. 1a). First, we designed a novel amplicon panel using our open-source multiplex PCR primer design software, *Multiply*^32^ (Methods). We targeted nine genes that would jointly provide information on antimalarial drug resistance, *hrp2/3* deletions, vaccine target evolution and complexity of infection (COI) (Table 1). Each gene was targeted with a single amplicon, except *mdr1* for which separate N- and C-terminal amplicons were designed. The amplicons vary in size from 604 bp to 1,405 bp and amplify a combined 10,860 bp in a single reaction. A total of 45 validated and candidate World Health Organisation (WHO) antimalarial drug resistance-associated single-nucleotide polymorphisms (SNPs) are genotyped (Table 1). The amplicons targeting *hrp2* and *hrp3* both have reverse primers annealing in exon 1, and forward primers upstream of exon 2, which enables detection of partial or complete hrp gene deletions^33^. The *csp* amplicon spans from codon 19 to the end of the gene, which includes the entire central repeat region and C-terminal domain used in the RTS,S/AS01 and R21/Matrix-M vaccines^34,35^. A high diversity, 826 bp window of *apical membrane antigen 1* (*ama1*) is covered to support COI estimation. We named this amplicon panel the NOMADS Minimal Viable Panel (MVP).

**Table 1.** Amplicons included in NOMADS-Minimal Viable Panel (MVP). For *kelch13* all WHO candidate and validate markers are spanned^36^. *mdr1*(n) and *mdr1*(c) indicate N- and C-terminal *mdr1* amplicons respectively. bp, base pairs. ^†^Amplicon spans C-terminal codon.

| No. | Reference ID | Gene Name | Amplicon (bp) | Codons Spanned | Key Regions |
| --- | --- | --- | --- | --- | --- |
| 1 | PF3D7_1133400 | *ama1* | 826 | 74 – 384 |  |
| 2 | PF3D7_0709000 | *crt* | 692 | 14 – 125 | K76T |
| 3 | PF3D7_0304600 | *csp* | 1,273 | 19 – 398<sup>†</sup> | RTS,S and R21 epitopes |
| 4 | PF3D7_0417200 | *dhfr* | 1,405 | 1 – 410 | A16V, N51I, C59R, S108M, I164L |
| 5 | PF3D7_0810800 | *dhps* | 1,327 | 317 – 707<sup>†</sup> | S436A/F, A437G, K540E, A581G, A613T/S |
| 6 | PF3D7_0831800 | *hrp2* | 1,296 | 14 – 306<sup>†</sup> |  |
| 7 | PF3D7_1372200 | *hrp3* | 1,216 | 14 – 276<sup>†</sup> |  |
| 8 | PF3D7_1343700 | *kelch13* | 1,286 | 383 – 727<sup>†</sup> | BTB/Propeller |
| 9 | PF3D7_0523000 | *mdr1*(n) | 935 | 46 – 245 | N86Y/F, Y184F |
| 10 | PF3D7_0523000 | *mdr1*(c) | 604 | 968 – 1278 | S1034C, N1024D, D1246Y |

**Figure 1.**
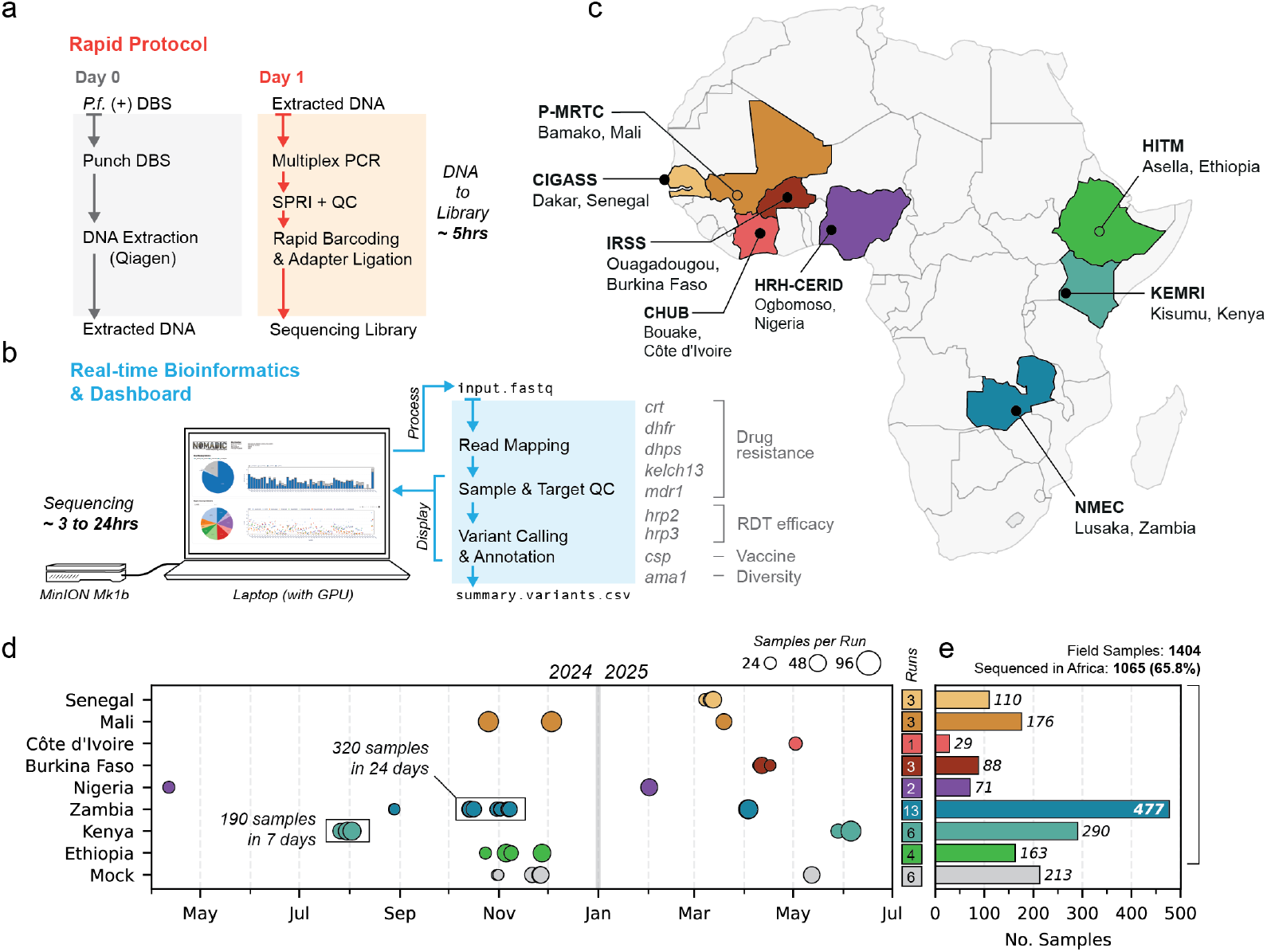
Overview of sequencing approach and implementation across sub-Saharan Africa. (a) The laboratory protocol uses *P. falciparum* positive dried-blood spots (DBS) as source material for DNA extraction. From extracted DNA to sequencing takes approximately 5 hours. (b) Data analysis occurs in real-time on a laptop. While sequencing is ongoing, quality control (QC) and variant calling results are displayed to an interactive dashboard. GPU: graphics processor unit. Sequencing time depends on flow cell quality and number of samples. (c) DBS samples were processed from 8 countries across sub-Saharan Africa. For six countries, all sequencing occurred locally (filled circles); for two countries, samples were sequenced internationally (Mali, Ethiopia; open circles). (d) Timeline of sequencing runs. Size of point indicates number of samples per run. Total sequencing runs for each country is shown in a box at right. (e) Barplot displaying total number of samples sequenced per country. Overall, 1,065/1,404 (65.8%) of samples were processed in Africa.

To promote widespread adoption, we developed a nanopore sequencing protocol for the NOMADS-MVP amplicons that minimises costs, complexity, and time. We initially explored two versions of the protocol: one that included a selective whole genome amplification (sWGA)^37^ step to preamplify bulk *P. falciparum* DNA, and one where the multiplex PCR was performed directly on extracted DNA. We reduced the run time of the sWGA step from 17 hours^37^ to 1 hour while still providing robust yield by using the EquiPhi29 polymerase (Methods). We hypothesised that including sWGA might increase sensitivity, although at additional protocol cost, complexity and time. While this was confirmed with mock DBS samples created from laboratory strains, we saw no benefit on field DBS samples and discontinued use of sWGA in August 2024. We optimised the NOMADS-MVP multiplex PCR using gradient PCR and several full factorial experimental designs: varying template DNA, primer and polymerase concentrations, annealing temperature, and the number of thermal cycles (Supplementary Fig. 1). After optimisation, multiplex PCR conditions maximised sensitivity (with amplification on mock DBS samples down to 100 parasites/µL) and simplicity. The final PCR only requires pipetting three reagents (primers, KAPA HiFi ReadyMix buffer, and sample), thus reducing contamination risk and increasing consistency. Finally, we adopted a rapid barcoding approach that dramatically reduces library preparation time and complexity, and simplifies procurement as no additional third-party reagents are required.

Starting from extracted DNA, the final optimised protocol costs $21 USD/sample, assuming 48 samples are sequenced in a batch and including all plasticware ($5.47 USD/sample, 26.2%) and reagents ($15.45 USD/sample, 78.3%; Supplementary Table 1). The three most costly items are the flow cells ($7.2 USD/sample, 34.4%), the KAPA HiFi polymerase ($3.19 USD/sample, 15.3%) and the 20 *µ*L multichannel pipette tips ($2.50 USD/sample, 12.0%). All the required equipment (including a laptop suitable for analysis) costs an estimated $12,500 USD (Supplementary Table 2). To objectively measure protocol complexity, we quantified the number of pipetting steps from extracted DNA to sequencing (1 step = aspirating and dispensing a reagent or sample). For a batch of 48 samples, our protocol requires 172 pipetting steps, compared to 292 steps for our previous long-read protocol^32^ or over 400 steps for two state-of-the-art Illumina-based protocols^23,31^ (Supplementary Table 3). The total incubation time for all steps in the NOMADS-MVP protocol is approximately 3.5 hours. After gaining familiarity, the workflow from extracted DNA to sequencing takes approximately 5 hours (Fig. 1a, Supplementary Fig. 2).

### A real-time, point-of-use bioinformatics pipeline and dashboard

We developed a real-time, on-site bioinformatics pipeline and analysis dashboard which accompanies the NOMADS-MVP protocol, called *Nomadic* (Fig. 1b). *Nomadic* maps reads, computes quality control statistics for each sample and amplicon, performs preliminary variant calling and annotation, and presents this information in a graphical dashboard while sequencing is ongoing. It allows laboratory scientists to make informed decisions regarding sample quality and amplicon performance, determine when to stop sequencing, and provides an immediate first-look at mutations present across samples. The dashboard is easy to install and is launched with a single command from the terminal, enabling use by scientists without significant bioinformatics training. Beyond what is required to perform real-time basecalling with *MinKNOW* (a laptop with a CUDA-enabled graphics processor unit), *Nomadic* requires no additional computational infrastructure and runs offline. In addition, *Nomadic* produces human-readable summary files that contain all key information about sequencing performance and detected mutations. These files are small enough (<20 Mbp) to be easily shared even over slow internet connections and are sufficient to reopen the graphical dashboard after the experiment is completed. *Nomadic* was used to process all the data described in this manuscript at the point of sequencing. Further details on *Nomadic* use and implementation are provided in the Methods.

### Implementation at scale across sub-Saharan Africa

From April 2024 to July 2025, a total of 1,404 dried-blood spot (DBS) samples from eight countries across sub-Saharan Africa were sequenced using the NOMADS-MVP protocol (Fig. 1c-e). Of these, 1,065 (65.8%) were sequenced locally, with the protocol being established in six countries: Nigeria (April 2024), Kenya (July 2024), Zambia (August 2024), Senegal (March 2025), Burkina Faso (April 2025), and Côte d’Ivoire (May 2025) (Fig. 1c). Overall, 13 local scientists independently conducted 28 sequencing runs across these six countries (Fig. 1d). The NOMADS-MVP protocol enabled periods of high sample processing throughput. For example, in Kenya two scientists processed 190 samples in 7 days at the end of July 2024 (Fig. 1d); and in late 2024, 320 samples were processed in 24 days by two scientists in Zambia.

Implementation of the protocol was feasible across laboratories that varied considerably with respect to prior sequencing experience and available infrastructure. One team had both Illumina and ONT sequencing experience (CIGASS, Senegal); three teams had previous nanopore sequencing experience (NMEC, Zambia; HRH-CERID, Nigeria; CHUB, Côte d’Ivoire); and for the remainder our protocol was the first experience with NGS. Infrastructure ranged from well-equipped regional sequencing hubs (CIGASS, Senegal) to small container laboratories with only the essentials for nanopore sequencing (NMEC, Zambia). For five countries, an initial in-person training was conducted by a core team (typically 1–3 weeks in duration) and followed by remote support. In Côte d’Ivoire, the protocol was established locally through a collaboration with the Robert Koch Institute (Berlin, Germany). Overall, these results demonstrate that our approach is amenable to large-scale implementation across a wide range of contexts in sub-Saharan Africa.

### Robust sequencing coverage across countries and parasitemia levels

We examined the sequencing coverage generated by the NOMADS-MVP protocol on a set of 1,283 samples, comprising 1,154 field DBS samples (1,154/1,283, 89.9%) and 129 mock DBS samples (129/1,283, 10.1%), all processed without sWGA (Fig. 2). Parasitemia data were available for 950 samples (950/1,283, 74.0%), which together had a median of 3,645 parasites/µL (IQR 631–11,540 parasites/µL). To evaluate sequencing performance, we calculated two per-sample metrics: (1) the mean coverage across all amplicons; (2) the fold-difference in coverage between the most- and least-abundant amplicon (excluding the *hrp2/3* amplicons), a measure of uniformity where a value of 1 indicates perfectly uniform coverage across amplicons (see Methods).

Across all samples, the median per-sample mean coverage was 1,881× (IQR 1,064–3,086×), with 76.3% (881/1,154) of field DBS samples and 87.5% (113/129) mock DBS samples achieving a mean coverage of greater than 1,000×. Across countries, the median per-sample mean coverage ranged from 544× in Côte d’Ivoire (*n* = 29) to 4,161× in Burkina Faso (*n* = 88), and 6 of 8 countries had a median per-sample mean coverage exceeding 1,000× (Fig. 2a). The median fold-difference in coverage across all samples was 10.2× (IQR 4.8–53.6×), with samples sequenced in Senegal having the most uniform coverage across amplicons (median 2.97×, IQR 2.14–4.95*×, n* = 110) and those from Zambia had the least uniform coverage (median 44.4×, IQR 9.5–197.4*×, n* = 457).

**Figure 2.**
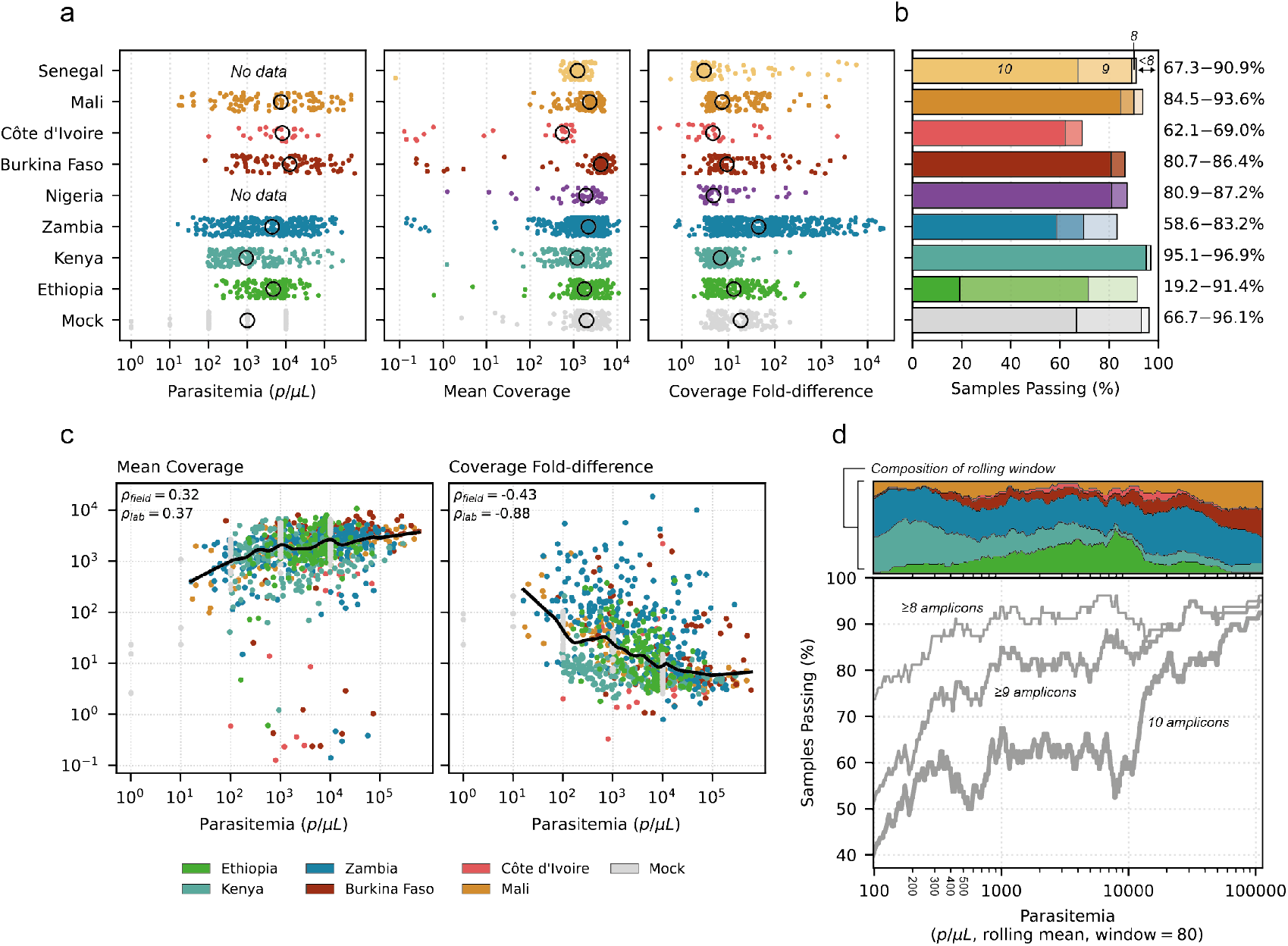
Sequencing coverage across countries and parasitemia levels. (a) Strip plots of parasitemia and sequencing coverage data for 1,283 samples processed with the NOMADS-MVP protocol, grouped by country. Each point represents a sample and black circles indicate country-level medians. The left subpanel shows sample parasitemia (parasites/µL); the middle subpanel shows mean per-sample coverage across amplicons; and the right subpanel shows per-sample coverage fold-difference between the least and most abundant amplicon (see Methods). All x-axes are on a logarithmic scale. (b) Bar plot showing, for each country, the percentage of samples with > 50 × coverage for 8 amplicons (lightest shade), 9 amplicons (intermediate), or all 10 amplicons (darkest); bar height indicates the total percentage of samples with > 50 × coverage for ≥ 8 amplicons. At right, percentages for 10 amplicons and ≥ 8 amplicons passing > 50 × coverage is annotated. (c) Scatter plots of parasitemia against mean coverage (left subpanel); and coverage fold-difference (right subpanel). Each point is a sample, colored by country. Black lines are a locally weighted scatterplot smoothing (LOWESS). Spearman’s *ρ* for the field (*ρ*_*field*_) and lab (*ρ*_*lab*_) samples is annotated at top left. Both axes use logarithmic scales. (d) Relationship between parasitemia and samples passing (%). Field samples were ordered by parasitemia, and rolling means were computed using windows of 80 samples. Lines represent percentage of samples with > 50 × coverage for ≥ 8 amplicons (thinnest line), ≥ 9 amplicons (medium), and for all 10 amplicons (thickest). The top subpanel shows the proportion of samples from each country (by colour) across the rolling parasitemia windows. Only field samples were included.

Next, for each sample we quantified the number of amplicons with greater than 50× coverage, which is sufficient for high-accuracy variant calling. Overall, 96.1% (124/129) of mock DBS samples and 87.9% (1,015/1,154) of field DBS samples had at least 8 amplicons with greater than 50× coverage. In 68.2% (88/129) of mock DBS samples and 64.5% of field DBS samples (745/1,154) all 10 amplicons exceeded 50× coverage. By country, the percentage of samples with ≥ 8 amplicons exceeding 50× coverage ranged from 69% in Côte d’Ivoire to 95% in Kenya (Fig. 2b). Combined, these results demonstrate that NOMADS-MVP can generate robust sequencing coverage across a diversity of sample sets and laboratories in sub-Saharan Africa.

Two major factors influencing sequencing performance are DNA quality and sample parasitemia. We observed a moderate positive correlation between parasitemia and per-sample mean coverage for both field DBS samples (Spearman’s *ρ* = 0.32, *p <* 0.001) and mock DBS samples (*ρ* = 0.37, *p <* 0.001). For example, in field DBS samples, the mean coverage at 1,000 parasites/µL (2,099×) was about twice that at 100 parasites/µL (1,011×). A stronger, negative correlation was observed between parasitemia and the per-sample fold-difference in coverage across amplicons (field DBS: *ρ* = *−*0.43, *p <* 0.001; mock DBS: *ρ* = *−*0.88, *p <* 0.001); this increased imbalance in coverage at lower parasitemia levels was driven primarily by decline (and occasional drop-out) of the two longest amplicons, which target *dhps* and *dhfr* (Supplementary Fig. 3). Finally, we evaluated the percentage of field DBS samples that had 8, 9, or 10 amplicons with greater than 50× coverage as a function of parasitemia. These percentages were consistently high for samples with *≥*1,000 parasites/µL, declining steadily below this threshold. For example, at 1,000 parasites/µL greater than 90% of samples had at least 8 amplicons with 50× coverage, compared to approximately 75% of samples at 100 parasites/µL. Including field DBS samples from Ethiopia, some of which had *hrp2/3* deletions, resulted in a lower percentage of samples with 9 or 10 amplicons exceeding 50× coverage. The mock

DBS samples generated from laboratory strains performed consistently well even at 100 parasites/µL (Supplementary Fig. 4). One explanation is that while mock DBS samples recapitulate absolute parasitemia, they represent a best-case scenario in terms of DNA quality, which may be challenging to achieve in the field.

### Rapid sequencing produces sufficiently long reads

The Rapid Barcoding Kit (SQK-RBK114.96) from ONT leverages transposome-based chemistry that cleaves sample DNA during barcode ligation. The barcoding reaction takes just four minutes, but the sample DNA is fragmented in the process. To characterise the impact on read lengths, we randomly selected 80 samples from across four different sequencing experiments, including field DBS samples from Kenya (*n* = 20), Ethiopia (*n* = 20) and Mali (*n* = 20), as well as a set of mock DBS samples created from laboratory strains in Germany (*n* = 20). Across these samples, a total of 3.45 million reads mapped uniquely to the *P. falciparum* reference genome and their median length was 567 bp (IQR 376–808 bp). There was considerably more read length variation within individual samples than between samples or experiments (Fig. 3a). The consistency across four independent sample sets and experiments suggests that the read length distribution is robust to technical and biological factors. Similarly, in optimisation experiments we found that read lengths were insensitive to variation in the barcoding reaction time or input DNA mass (Supplementary Fig. 5). The within-sample read length variation was driven by the different amplicons in our panel, with, as expected, longer amplicons generating longer reads (Fig. 3b). In particular, when grouped by the amplicon to which they mapped, reads had a median length of roughly half of their amplicon length (mean 59%, range 45%–69%), consistent with a single cleavage and barcoding event per amplicon (Fig. 3b). Across amplicons, the longest median read length was for the *kelch13* amplicon at 872 bp and the shortest for the *mdr1* N-terminal amplicon at 397 bp.

**Figure 3.**
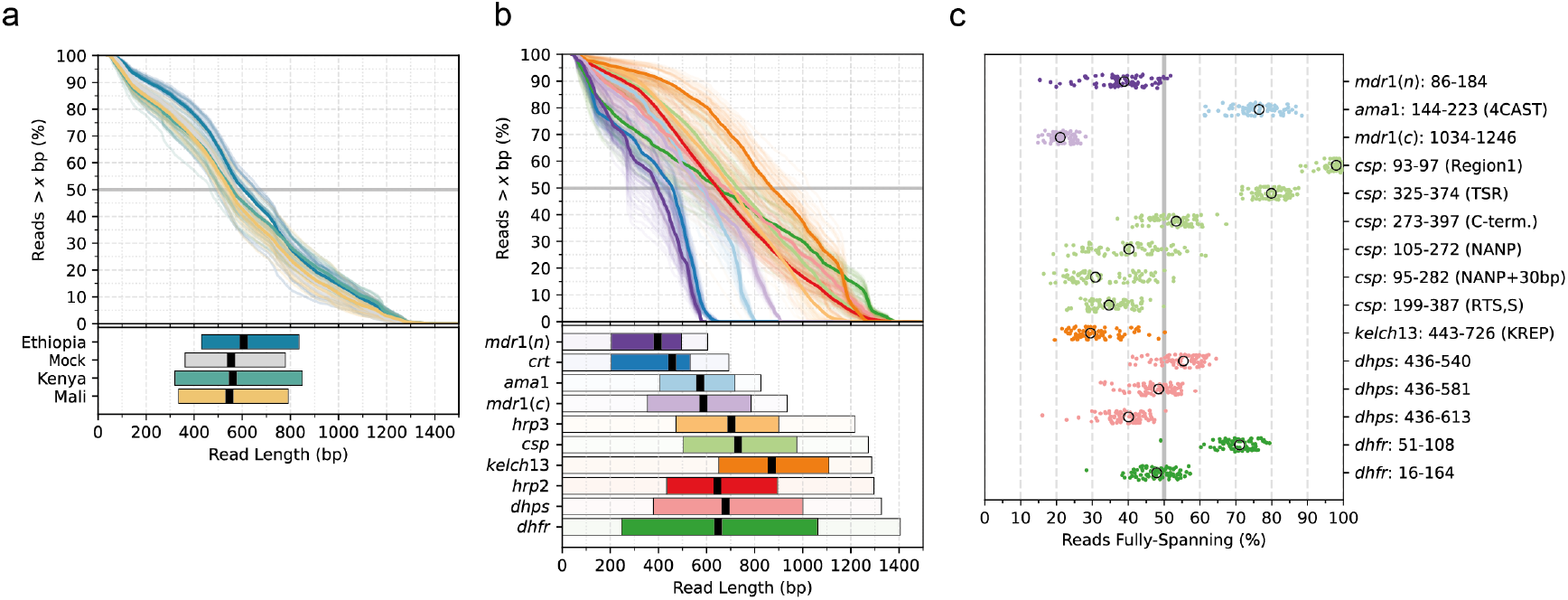
Read lengths generated by the NOMADS-MVP protocol. (a) Read length distributions for 80 samples from four countries. The top subpanel shows the percentage of reads (y-axis) longer than a given length (x-axis). Each line represents one sample; colours indicate country. The bottom subpanel shows box plots of read length distributions by country. Boxes indicate the interquartile range, black lines mark the median. (b) Same as (a), but reads are grouped by target amplicon rather by than sample. In the bottom subpanel, the transparent bar behind each box indicates the full amplicon length. (c) Strip plot showing the percentage of sequencing reads fully spanning a given genomic region of interest for each sample. Genomic regions are defined by gene and codon range. 4CAST, region of *ama1* amplified in *La Verriere et al*.^22^; TSR, thrombospondin-like type I repeat; KREP, Kelch-repeat propeller. Black circles indicate the region-level medians.

Longer reads support physical phasing of heterozygous variants and the investigation of repetitive regions, but only when the variants or regions are fully spanned by individual reads. To this end, we examined whether our data contained reads that fully spanned regions of interest for *P. falciparum* genomic surveillance (Fig. 3c). All regions investigated had fully-spanning reads at appreciable frequencies across the eighty samples. For example, a median of 29.4% (IQR 26.4%–33.5%) of reads per sample overlapping the Kelch-repeat propeller domain fully spanned the region. For *dhps*, which mediates Sulfadoxine resistance, a median of 40.0% (IQR 35.9–43.0%) of reads spanned from codon 436 to 613, enabling joint interrogation of the key resistance markers (S436A, A437G, K540E, A581G and A613S). A high-diversity region of *ama1*^22^ was spanned by a median of 76.5% (IQR 72.1–80.6%) of reads per sample. Finally, key regions of the RTS,S/AS01 and R21/Matrix-M vaccine target *csp* also had fully-spanning reads, including the entire portion used in the vaccine (codons 199–397, median 34.6% of reads fully-spanning) and the entire NANP central repeat region (codons 105–272, median 40.2%).

### A novel variant caller for polyclonal *P. falciparum* infections

Sensitive and specific detection of single-nucleotide polymorphisms (SNPs) are critical for accurate antimalarial drug resistance prediction from known genetic markers. However, in polyclonal *P. falciparum* infections, SNPs carried by low frequency minor clones (e.g. *<* 20%) are often undetected by diploid variant callers; including the main variant callers supporting nanopore sequencing data, such as *bcftools*^38^, *Clair3*^39^, *PEPPER-Margin-DeepVariant*^40^, or *Longshot*^41^. As a result, pipelines using these variant callers can have reduced sensitivity in polyclonal *P. falciparum* infections.

To address this issue, we developed a novel variant caller, named *Delve*, which enables sensitive SNP detection in polyclonal *P. falciparum* infections with low frequency minor clones. In brief, *Delve* calls biallelic SNPs with a statistical model that utilises the aligned bases and quality scores at each genomic position after read mapping (Methods). It removes the assumption of a diploid organism by reparameterising the genotype likelihood function used by *bcftools*^38^; expressing the likelihood in terms of a continuous within-sample alternative allele frequency (WSAF). The WSAF is free to take any value between zero and one, reflecting the fraction of minor clones that carry the alternative allele. *Delve* determines the maximum likelihood WSAF by numerical optimisation, and then assigns a genotype of either homozygous reference, heterozygous, or homozygous alternative using a series of likelihood ratio tests. Candidate SNPs are filtered based on quality and probable sequencing bias to generate the final SNP calls for the sample. We implemented *Delve* in Rust and it is open-source and publicly available on GitHub (Methods).

### Sensitive and precise SNP calling in polyclonal infections

To assess SNP calling performance, we created 45 mock DBS samples from the *P. falciparum* laboratory strains 3D7, Dd2 and HB3. These included clonal samples (3D7, Dd2, or HB3) and two-strain mixtures (3D7 and Dd2, 3D7 and HB3, or HB3 and Dd2) at different minor clone proportions (20%, 10%, 5% and 2.5%) and parasitemia levels (10,000, 1,000, and 100 parasites/µL). We extracted DNA from a single 6 mm punch of each mock DBS sample using a Qiagen spin-column protocol (Methods). We then performed the NOMADS-MVP protocol in triplicate, sequencing the 45 mock samples (plus 3 negative controls) thrice on three separate R10.4.1 flow cells (Fig. 4a). Each run produced a high mean coverage across all amplicons (run 1: 2,021×; run 2: 1,459×; run 3: 4,798×) which declined with parasitemia (3,529× at 10,000 parasites/µL versus 1,822× at 100 parasites/µL, averaged across runs); consistent with observations from field samples.

**Figure 4.**
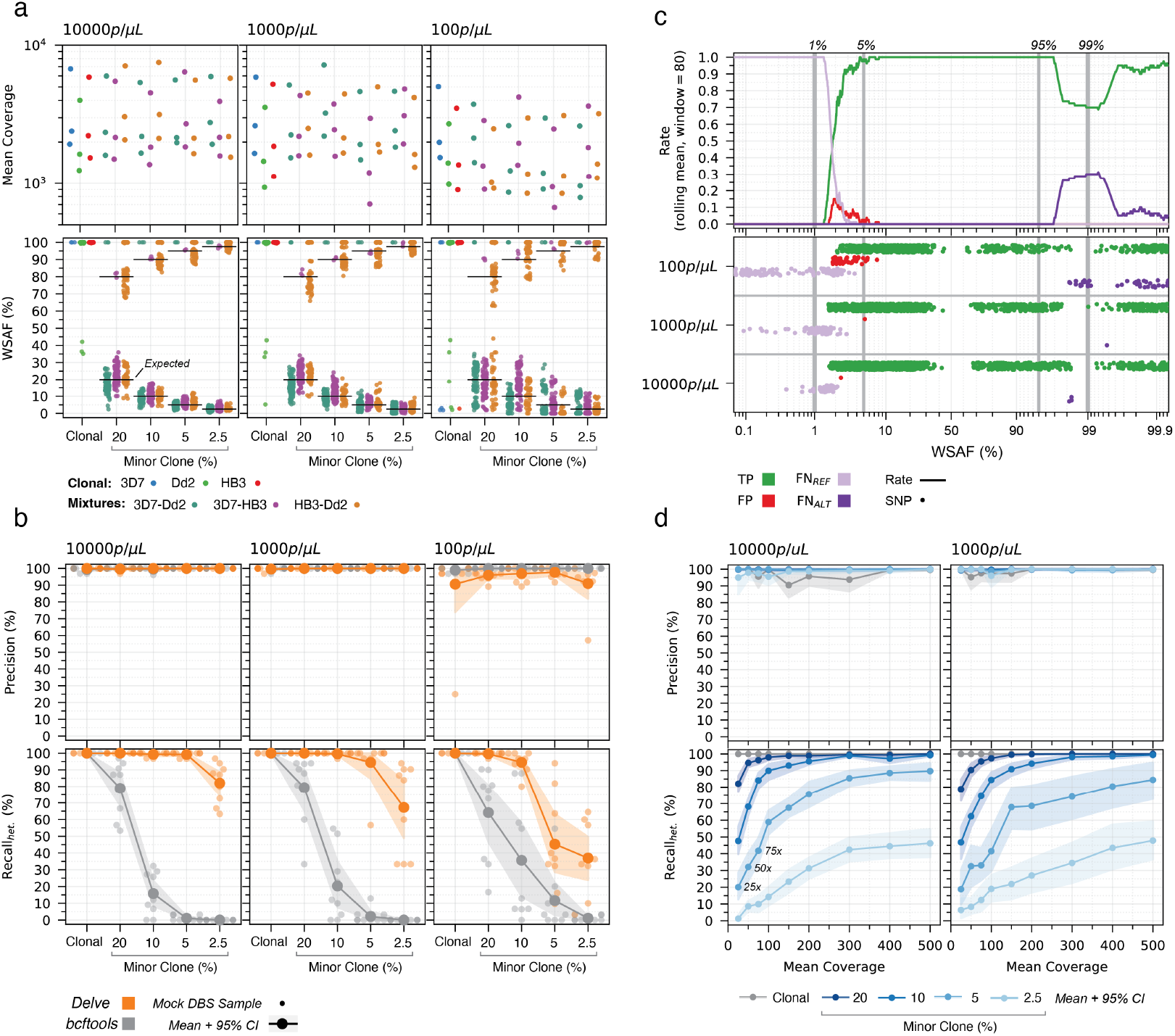
SNP calling performance in clonal and polyclonal mock DBS samples. A total of 45 mock DBS samples were sequenced in triplicate. (a) Strip plots showing the mean coverage per mock DBS sample (top), and WSAF for every observed SNP (bottom). Mock DBS samples are defined by their constituent strain(s) (color) and minor clone proportion (x-axis). In the bottom subpanels, the horizontal bars indicate the expected WSAF given the minor clone proportion. Note that due to duplication of *mdr1*, the clonal Dd2 samples have one heterozygous site. (b) Same arrangement as (a), but the top row of subpanels show the SNP calling precision, and the bottom show the recall of true heterozygous SNPs (Recall_*het*_), for both *Delve* (orange) and *bcftools* (grey). Small points represent individual mock DBS samples; the line and shaded area represent the mean and 95% confidence intervals, calculated by bootstrapping. The recall of homozygous alternative SNPs was perfect for both tools (data not shown). (c) Relationship between observed WSAF and SNP calling outcome for *Delve*. Each true or called SNP was classified as either a true positive (TP, green), false positive (FP, red), or false negative in which a heterozygous SNP was called homozygous reference (FN_*REF*_, light purple) or homozygous alternative (FN_*ALT*_, dark purple). SNPs were ordered by WSAF, and a rolling rate for each SNP category was computed using windows of 80 SNPs (top subpanel). Individual SNPs are shown in a strip plot (bottom subpanel) grouped by mock DBS sample parasitemia (y-axis). The majority of false-positive SNPs have a WSAF of *<*5%. (d) Effect of downsampling on precision (top) and the recall of heteryzgous SNPs (bottom). Mock DBS samples with 10,000 and 1,000 parasites/µL were downsampled in triplicate to different coverage levels (x-axis). Color indicates clonal (grey) or minor clone proportion (shades of blue). Lines and shaded areas represent the mean and bootstrapped 95% confidence intervals.

To confirm that the mock DBS samples were created accurately, we compared the observed within-sample alternative allele frequencies (WSAF, the percentage of reads carrying the alternative allele) for all true heterozygous SNPs against their expected values, which are derived from the minor clone proportion (Fig. 4a). Overall, the WSAF biases for heterozygous SNPs were small. For example, mock DBS samples with a minor clone at 10% had a median WSAF of 10.3% at 10,000 parasites/µL (IQR 8.0–12.5%, *n* = 231 heterozygous SNPs) and 10.3% at 1,000 parasites/µL (IQR 8.0–13.6%, *n* = 231); while mock DBS samples with a minor clone at 5% had a median WSAF of 5.6% at 10,000 parasites/µL (IQR 4.2–6.7%, *n* = 231) and 5.7% at 1,000 parasites/µL (IQR 4.0–6.9%, *n* = 231). These results confirmed that the mock DBS samples carried minor clones at the intended proportions and were suitable for interrogating the minor clone limit-of-detection (LoD).

The observed median WSAF across heterozygous SNPs reflects the mock DBS minor clone proportion, while variation around the expected WSAF is introduced primarily by molecular variability during DBS processing (i.e. DNA extraction, multiplex PCR, and library preparation), as well as potential sequencing error. Consistent with this, at 100 parasites/µL we observed greater WSAF deviations than at higher parasitemia levels (100 parasites/µL: root mean squared error [RMSE] 7.0%, *n* = 1, 127 heterozygous SNPs; 1,000 parasites/µL: RMSE 4.0%, *n* = 1, 128; 10,000 parasites/µL: RMSE 3.3%, *n* = 1, 127). Moreover, we found that haplotypes carried by the 5% and 2.5% minor clones were often absent at 100 parasites/µL: in 39.6% (25/63 amplicons) and 55.0% (33/60 amplicons) of cases, respectively, they produced no WSAF signal (Supplementary Fig. 6). However, effectively all haplotypes with a minor clone proportion of 10% at 100 parasites/µL generated signal (98.3%, 59/60 amplicons). This suggests that, regardless of the subsequent bioinformatics analysis, the NOMADS-MVP protocol has a minimum minor clone LoD of above approximately 5 parasites/µL (5% of 100 parasites/µL), set by the molecular stochasticity and the sensitivity of the laboratory steps. Achieving a minor clone LoD below this would require, for example, using more efficient DNA extraction protocols or developing more sensitive PCRs.

Next, we called SNPs using both *Delve* and *bcftools*. We compared their results to the true SNPs for each mock DBS sample and evaluated performance by calculating precision and recall (Fig. 4b). To isolate minor clone detection performance, we calculated separate recall values for the true homozygous alternative SNPs and true heterozygous SNPs. The mean recall of homozygous alternative SNPs was perfect (100%) for both tools. *Delve* had a higher recall of heterozygous SNPs than *bcftools* in all 108 polyclonal mock DBS samples (Fig. 4b). In samples with minor clones at 10%, the recall of heterozygous SNPs was high at all parasitemia levels with *Delve* (10,000 parasites/µL: 99.3%; 1,000 parasites/µL: 99.6%; 100 parasites/µL: 94.5%) but low with *bcftools* (10,000 parasites/µL: 15.8% *bcftools*; 1,000 parasites/µL: 20.3%; 100 parasites/µL: 35.6%). Overall, the data indicated that the approximate LoD for minor clones when using NOMADS-MVP and calling SNPs with *Delve* is 5% at 10,000 parasites/µL (mean recall 99.2%, *n* = 9 samples), 5% at 1,000 parasites/µL (mean recall 94.4%, *n* = 9) and 10% at 100 parasites/µL (mean recall 94.5%, *n* = 9).

We examined the false-positive and false-negative errors in more detail. The mean precision across all mock DBS samples and replicates with 1,000 parasites/µL or greater (*n* = 90) was very high: 99.9% for *Delve* and 99.8% for *bcftools*. At 100 parasites/µL, however, the mean precision of *Delve* was lower than *bcftools* (94.5% versus 99.7%, *n* = 45 samples), with *Delve* calling a mean of 0.95 false-positive SNPs per sample (range 0–4) at this parasitemia level. All of the false-positives had a low WSAF (median 2.8%, IQR 2.1–3.9%, *n* = 45), with 88% (40/45) having a WSAF less than 5% (Fig. 4c). In addition, the false-positives had much weaker statistical evidence in support of a SNP (likelihood ratio test [LRT] statistic, median 24.5, IQR 13.4–43.9) than did true-positive heterozygous SNPs (LRT statistic, median 606.1, IQR 156.4–2,266.4); it would, therefore, be possible to filter them, but with a penalty to recall. For all false-negatives in which a true heterozygous SNP was incorrectly called homozygous reference, the WSAFs were also very low (median 0.1%, IQR 0.0–1.3%, max 3.7%, *n* = 422); conversely, for false-negatives in which a true heterozygous SNP was called as homozygous alternative, the WSAFs were very high (median 99.9%, IQR 99.1–100%, min 98.1%, *n* = 80; Fig. 4c). This suggests that the majority of false-negatives are caused by molecular stochasticity and sensitivity of the NOMADS-MVP protocol (as discussed above), rather than by *Delve*. Significantly, across all mock DBS samples (including those at 100 parasites/µL), when the WSAF was between 5 and 95%, *Delve* detected all heterozygous SNPs and had a precision of 99.7% (2195 of 2200 called SNPs were true-positives).

Finally, we used downsampling to explore the relationship between sequencing coverage and the SNP calling performance of *Delve* (Fig. 4d). We randomly sampled sequencing reads to achieve a specific coverage ranging from 500 *−* 25×, in triplicate, for all mock DBS samples with 1,000 parasites/µL or greater and sufficient coverage across all amplicons (Methods). Similar to with the non-downsampled mock DBS samples, the recall of true homozygous alternative SNPs was perfect (100%, *n* = 1911 samples) and the mean precision was very high (99.1%, mean of 0.07 false-positive SNPs per sample, range 0–2, *n* = 1, 911 samples) for all samples in this analysis. The few examples where mean precision noticeably declined in Figure 4d were caused by the clonal 3D7 mock DBS samples (e.g. 150× mean coverage at 10,000 parasites/µL: 93.4% including 3D7 [*n* = 27] versus 99.6% excluding 3D7 [*n* = 18]); 3D7 has only 1 true positive SNP, and so a single false-positive reduces the precision to 50% and substantially depresses the mean. For mock DBS samples with 20% minor clones, *Delve* had a high recall of heterozygous SNPs with only 75× coverage (10,000 parasites/µL: 96.3%, *n* = 21 samples; 1,000 parasites/µL: 95.4%, *n* = 21 samples). For mock DBS samples with 10% minor clones, *Delve* had a high recall of heterozygous SNPs with 200× coverage (10,000 parasites/µL: 95.5%, *n* = 21 samples; 1,000 parasites/µL: 94.2%, *n* = 21 samples). For mock DBS samples with 5% minor clones, *Delve* achieved a mean recall of 89.6% at 10,000 parasites/µL and 84.4% at 1,000 parasites/µL with 500× coverage. For context, we compared these coverage levels to those achieved for field DBS samples sequenced in this study. In all field DBS samples passing quality control, 95.4% (10,886/11,390) of amplicons exceeded 75× coverage, 91.4% (10,408/11,390) exceeded 200× coverage, and 82.1% (9,362/11,390) exceeded 500× coverage. In summary, this analysis suggests that NOMADS-MVP and *Delve* should enable routine detection of minor clones at frequencies down to between 5 and 10% in the field.

### Accurate identification of antimalarial drug resistance-associated mutations

Antimalarial resistance marker identification is central to genomic surveillance. Therefore, we specifically investigated the recovery of WHO-defined antimalarial resistance markers^9^ across the mock DBS samples described in the section above; as well as in mock samples created from three Cambodian *P. falciparum* strains carrying validated artemisinin resistance mutations (*kelch13* R539T, I543T, and C580Y), and sequenced in duplicate. In clonal mock samples (*n* = 33), *Delve* identified all antimalarial resistance markers with 100% accuracy (no false-positives or false-negatives; Supplementary Table 4). *Delve* also identified all resistance markers with 100% accuracy in polyclonal mock DBS samples with 1000 parasites/µL or greater down to a minor clone frequency of 5%, except for *mdr1* N86Y (accuracy 77.8%; Supplementary Table 4). Because the *mdr1* gene is present in two or three copies in our *Dd2* clone, the N86Y marker WSAF is typically less than half the minor clone frequency, making the mutation harder to detect and explaining the reduced accuracy. At 100 parasites/µL, *Delve* identified antimalarial resistance markers with 98.8% and 92.2% accuracy for polyclonal mock DBS samples with minor clones at 20% and 10% frequency, respectively (Supplementary Table 4). Importantly, no false-positive SNP impacted any codon associated with antimalarial drug resistance; the observed reductions in accuracy were exclusively due to false-negatives arising when markers carried by minor clones fell below the LoD.

### Reliable and streamlined detection of *hrp2/3* deletions

We evaluated the ability of NOMADS-MVP to detect *hrp2* and *hrp3* deletions, which can cause false-negative RDT results and are currently prevalent in the Horn of Africa^2,3,5,6^. We used a set of 149 DBS samples collected from malaria patients in central Ethiopia, which had been previously assayed for *hrp2/3* deletions using recommended conventional PCR assays performed in duplicate^42^. We sequenced the 149 samples across four MinION sequencing runs, including in each run mock DBS samples created from the laboratory strains 3D7 (*hrp2*+/*hrp3*+), Dd2 (*hrp2−*/*hrp3*+) and HB3 (*hrp2*+/*hrp3−*), at parasitemia levels ranging from 100–10,000 parasites/µL; as well as mock *P*.*falciparum*-negative samples. We inferred *hrp2/3* deletions using a statistical model that estimates sample quality (from the mean abundance of other amplicons in the panel) and the rate of background sequencing contamination (from the negative controls), before calculating a deletion probability for *hrp2* and *hrp3*^32^.

NOMADS-MVP correctly estimated the *hrp2/3* deletion status for all mock samples (Fig. 5a). Mean amplicon coverage tended to decline with parasitemia and low levels of sequencing contamination were evident in some *P. falciparum*-negative controls. Notwithstanding, for both *hrp2* and *hrp3*, the difference in coverage between mock samples with and without deletions consistently exceeded 100-fold. For example, the mean *hrp2* amplicon coverage in Dd2 versus 3D7 mock samples was 6.6× versus 2,340× at 10,000 parasites/µL; 1.8× versus 1,968× at 1,000 parasites/µL; and 1.1× versus 984× at 100 parasites/µL (Fig. 5b). Similarly, for *hrp3* the mean amplicon coverage in HB3 versus 3D7 was 8.8× versus 4,910× at 10,000 parasites/µL; 3.3× versus 3,716× at 1,000 parasites/µL; and 2.0× versus 2,262× at 100 parasites/µL (Fig. 5c).

Of the 149 field DBS samples from Ethiopia, 138 (92.6%) passed quality control. Among these, the joint *hrp2/3* deletion results from conventional PCR matched those from NOMADS-MVP in 136 samples (98.6%; Fig. 5d). The two discrepancies included one sample that was *hrp2−*/*hrp3−* by NOMADS-MVP but *hrp2*+/*hrp3−* by conventional PCR (Fig. 5c, indicated by ‘ii’). This sample had a mean coverage of 425× over the *hrp2* amplicon; a much higher mean coverage than other samples predicted as *hrp2−* (median 0.3×, IQR 0.0–0.9×), but lower than those predicted *hrp2*+ (median 1,468×, IQR 934–3,433). Heterozygosity over *ama1* indicated that the sample was a mixed infection with at least two *P*.*f*. clones. A possible explanation for this discrepancy is that one clone carries an *hrp2* deletion that is detected by the statistical model, but still yields a positive result by conventional PCR. Similarly, the second discrepancy (*hrp2−*/*hrp3−* by NOMADS-MVP, *hrp2−*/*hrp3*+ by conventional PCR) had non-negligible coverage over *hrp3* (78×) and *ama1* diversity suggesting a mixed infection (Fig. 5c, indicated by ‘iii’). We also noted one other sample where, although conventional PCR and NOMADS-MVP both predicted an *hrp3* deletion, considerable coverage over the *hrp3* (479×) was present (Fig. 5c, indicated by ‘i’). None of the discordant samples had particularly low parasitemia (i: 2,960 parasites/µL; ii: 4,960 parasites/µL; iii: 9,080 parasites/µL) or mean coverage (i: 1,505×; ii: 1,752×; iii: 3,753×), which might be expected if insufficient sensitivity was an underlying cause. All samples whose parasitemia or mean coverage was in the lowest decile (229–960 parasites/µL; or 323–647×) had corresponding *hrp2/3* deletion calls between the PCR-based assays and NOMADS-MVP. Overall, these results demonstrate that NOMADS-MVP is a robust method for the detection of *hrp2/3* deletions, yielding results that align closely with conventional PCR assays currently in widespread use. In addition, it is more streamlined than existing laboratory workflows^43^; generating results for both *hrp2* and *hrp3* simultaneously, and because the other amplicons act as internal controls, without requiring secondary PCRs to confirm sample DNA quality.

## Discussion

Genomic surveillance of *P. falciparum* malaria can generate valuable public health data, but implementation across sub-Saharan Africa has been limited by a lack of sequencing approaches suitable for most local laboratories. Here, we described the development of a novel nanopore sequencing protocol for *P. falciparum* genomic surveillance which targets ten genomic regions, collectively providing information on a panel of antimalarial drug resistance-associated genes, *hrp2/3* deletions, the vaccine target *csp*, and the highly diverse gene *ama1*. In one year, we implemented the protocol in Senegal, Burkina Faso, Côte d’Ivoire, Nigeria, Zambia and Kenya — locally sequencing and analysing 1,065 DBS samples — and demonstrating the feasibility of continental-scale decentralised *P. falciparum* genomic surveillance.

After genomic data are generated, difficulties performing bioinformatic analysis in sub-Saharan Africa often create another barrier to timely surveillance. We addressed this by developing a real-time bioinformatics dashboard called *Nomadic*, which has two main advantages. First, it can be run with minimal bioinformatics expertise, enabling a wide range of users to independently and immediately derive insights from their data. Second, it runs offline on the same laptop used for sequencing, making it affordable and invulnerable to the internet connectivity issues which can bottleneck workflows reliant on transferring data to remote servers^30^. While analysis occurs on-site and raw data can remain local, *Nomadic* produces compact summary files that can be easily shared. These summary files are useful for collaborative troubleshooting and, in the future, could facilitate regional data integration when decentralised genomic surveillance becomes widespread.

Identifying variants associated with antimalarial drug resistance is a core aim of genomic surveillance. Yet in polyclonal *P. falciparum* infections, variants carried by minor clones are often missed by standard variant callers, which (incorrectly) assume a diploid organism. We addressed this by developing a novel variant caller, named *Delve*, and demonstrated its ability to recall SNPs carried by minor clones down to 5% frequency while maintaining a high precision. We highlight that *Delve* is a relatively simple variant caller, calling only biallelic SNPs, and refraining from recalibrating base quality scores, sharing information across variant sites, or using prior information about population-level allele frequencies. It is possible that more complex somatic variant callers (e.g. *Strelka2*^44^) could be calibrated to *P. falciparum* nanopore sequencing data and outperform *Delve*, although this has not been explored. We also highlight that haplotype calling tools originally designed for Illumina data, such as *DADA2*^45^ and *SeekDeep*^46^, have been increasingly applied to nanopore data^30^ to detect low frequency minor clones. Haplotypes are more informative than SNPs, however, these haplotype calling tools require unfragmented reads derived from complete amplicons; making them incompatible with the transposase-based rapid barcoding kits (e.g. SQK-RBK114.96, ONT) that enable much faster and simpler laboratory protocols for nanopore sequencing. Rather than inferring haplotypes directly, the long-reads produced by NOMADS-MVP (Fig. 3) could be used to physically phase SNPs called by *Delve* with a tool like *WhatsHap*^47^. Many salient drug-resistance phenotypes depend on combinations of SNPs, and this approach will enable NOMADS-MVP to predict them accurately even in polyclonal *P. falciparum* infections.

Our protocol does have limitations that make it less well suited for some malaria genomic surveillance activities. The longer amplicons (500–1,500 bp) generated allow contiguous coverage of target genes and simpler multiplexing, but also require samples to have higher parasitemia and/or DNA quality for robust amplification. For clinical cases of malaria we have demonstrated robust performance with NOMADS-MVP across a wide range of studies. However, for asymptomatic or submicroscopic cases short-read sequencing approaches are likely to have more consistent performance and less dependence on DNA quality, although we note that several studies included here contained asymptomatic cases and performed well (including dry-season samples from Mali and community-collected samples from Zambia). Similarly, for studies distinguishing *P. falciparum* recrudescence from reinfection, short-read approaches may enable more sensitive minor clone detection^23,30,31^. Beyond sensitivity, our assay lacks some outputs generated by other assays, such as between sample relatedness estimates. Relatedness and identity-by-descent (IBD) are best estimated by leveraging a larger number of unlinked variant sites (10s to 1000s) spread across the genome. Highly multiplexed short-read amplicon sequencing^21–23^, MIPs^24,25^, SNP barcodes^48^ or whole-genome sequencing (WGS) methods are better suited to make these estimates for studies that require them. Accurate COI estimation will be possible leveraging the *ama1* and *csp* amplicons and work to provide this output is underway.

The limitations of NOMADS-MVP protocol are counterbalanced by its ability to detect important threats to malaria control and case management with simplicity and rapidity — thereby meeting an urgent public health need. It is feasible to sequence a moderate sized batch of samples (24–64) and use *Nomadic* to generate results within a single day, and with substantially less laboratory manipulation than other approaches. The protocol can be easily implemented even in small laboratories with limited infrastructure, internet connectivity, or prior sequencing experience. Because it takes only one or two days to complete, scientists both process sample sets faster and, in the context of training, gain mastery faster, repeating the protocol and accruing experience quickly. Reducing the set of required reagents makes procurement for the protocol easier, simplifies troubleshooting, and ultimately makes outputs more consistent. Moreover, with a rapid protocol the consequences of a disruption (e.g. by a power outage) are less profound, and scheduling sequencing around other activities is easier. Altogether, these features will accelerate decentralised *P. falciparum* genomic surveillance across sub-Saharan Africa — improving both the quantity and quality of data that are generated, while simultaneously strengthening local capacity and country ownership.

## Methods

### Sample collection and ethics

All blood samples were collected from patients with *P. falciparum* malaria, with informed consent from the patient or from a parent or guardian. In all cases, consent allowed for the samples to be used for purposes such as this study. This study was conducted using the collected samples only; there was no human subject contact.

All studies received ethical approval from an appropriate research ethics committee: in Senegal, ethical approval for the SEN19/49 study was granted by the Comité National d’Ethique pour la Recherche en Santé (000317/MSAS/CNERS/SP); in Torodo, Mali, ethical approval for a cohort study in 2022 was granted by the Ethics Committee of Charité (EA2/264/21) and the Faculty of Medicine, Pharmacy and Odontostomatology (FMPOS) at the University of Bamako (N°2022/20/CE/USTTB/24.01.22); in Burkina Faso, ethical approval for the AMTIP study was granted by the Comité d’Ethique Institutionnel / Institut National de Santé Publique (2023-10/MSHP/SG/INSP/CEI); in Côte d’Ivoire, ethical approval for the study “Surveillance génomique de Plasmodium falciparum aux CTA à Bouaké, Côte d’ivoire” was granted by the Direction Médicale et Scientifique du aux Centre Hospitalier et Universitaire (192MSHPCMU/CHU-B/DG/DMS/ONAR/24) and by the Ethikkomission der Charité Universitätsmedizin (EA2/171/24); in Nigeria, ethical approval for samples collected in 2023–2024 was granted by the Ethical Committee of Ladoke Akintola University of Technology Teaching Hospital, Ogbomoso (LTH/OGB/EC/2022/304); in Zambia, ethical approval for the 2024 Malaria Indicator Survey (MIS2024) and the 2024–2025 *hrp2/3* Surveillance Study was granted by the Research Ethics Committee at the University of Zambia (5055-20241) and the Tropical Diseases Research Centre Ethics Review Committee (TRC/C4//03/2024), respectively; in Kenya, ethical approval for samples collected in the ATSB study was granted by the KEMRI Scientific and Ethics Review Unit (KEMRI/SERU/CGHR/368/4189; CDC Project ID 0900f3eb81d7ec3c and 0900f3eb82546323); in Ethiopia, ethical approval for the ARSUNA study was granted by the Arsi University institutional review board (AU/HSC/ST-129/5494) and the Federal Democratic Republic of Ethiopia, Ministry of Education (17/256/476/24).

### Mock sample creation

#### Making DBS from culture

For validation of our protocol we created mock *P. falciparum* positive dried-blood spots (DBS) as follows. We cultured the *P. falciparum* laboratory strains 3D7, Dd2 and HB3 at 5% hematocrit in commercially available red blood cells (RBCs) obtained from DRK-Blutspendedienst Nord-Ost gGmbH, as previously described^49^. We brought each culture to ∼5% parasitemia and then synchronised to ring-stage parasites using 5% Sorbitol (PanReac AppliChem, #A2222). After synchronisation, three technicians independently measured the parasitemia of each culture by microscopy and used a Neubauer Counting Chamber to determine the number of RBCs per microlitre. We used the average of the parasitemia and RBCs/*µ*L measurements to standardise each culture to 100,000 parasites/µL at 50% hematocrit. We performed 10-fold serial dilutions of the 100,000 parasites/µL stocks in 80 *µ*L of whole human blood to produce clonal mock samples down to 1 parasites/µL. To explore sequencing performance on polyclonal infections with low frequency minor clones, we created mock samples containing two laboratory strains at different proportions. In particular, we combined clonal 10,000 parasites/µL dilutions of two strains to create 80:20 mixtures in 120 *µ*L, and then performed two-fold serial dilutions into the major strain to produce 90:10, 95:5, and 97.5:2.5 mixtures. These were further serial diluted into whole human blood to produce 1000 parasites/µL and 100 parasites/µL parasitemia mixtures. To produce the DBS, we created five 20 *µ*L spots for each mock sample on an individual filter paper (Whatman, #10531018) and dried them overnight at room temperature (Supplementary Fig. 4a). The DBS were stored at *−*20°C in individual plastic bags with a desiccant until use.

#### DNA extraction from DBS

For each sample we used a single (6mm) punch from the DBS for DNA extraction. DNA was extracted by using QIAamp DNA Mini Kit (Qiagen, #51306) according to the manufacturer’s instructions with the following modifications: elution was performed by using 2×40 *µ*L of nuclease free water heated to 50 °C. Each elution was incubated for 3 min at room temperature.

#### Mock samples containing kelch13 mutations

To interrogate artemisinin-resistance mutations, we ordered *P. falciparum* genomic DNA for Cambodian field derived strains IPC 5202 (*kelch13* R539T), IPC 4912 (*kelch13* I543T), IPC 3445 (*kelch13* C580Y)^50^ from BEI resources (www.beiresources.org). We created 10,000 parasites/µL *in vitro* DNA mixtures by diluting these stocks to 0.25 ng/*µ*L in 25 ng/*µ*L human genomic DNA (Roche #11691112001).

### NOMADS-MVP Multiplex PCR

#### Primer design

We designed primers for the NOMADS-MVP (Minimum Viable Panel) amplicon panel using our open-source software *Multiply* (https://github.com/JasonAHendry/multiply). In brief, *Multiply* takes as input a user-specified “design file” defining the target organism, regions of interest, acceptable amplicon size range, and other parameters. From the design file, it generates a large set of candidate primers for each target region and finds an optimal multiplex PCR primer set by minimising primer dimers, offtarget annealing, and primer overlap with known genetic variants. For moderate target numbers (> 20), it typically takes a few minutes to run. For the ten targets of NOMADS-MVP (Table 1), *Multiply* produced a total of 453 candidate primer pairs (median of 47 per target); across which it identified 4,096 potential offtarget binding sites, 77 primers overlapping common variants (>5% minor allele frequency (MAF) in any population defined in the Pf6 Project^51^ or resistance-associated codons), and 2,028 high-affinity primer dimers. The greedy search algorithm minimised these factors to select a set of 3 candidate multiplexes (from a theoretical 22 thousand trillion possible combinations) for the ten targets. After pilot sequencing experiments, we selected the multiplex with the highest coverage over all ten targets as our final panel (Supplementary Fig. 1d).

#### PCR optimisation

We optimised our multiplex PCR using KAPA HiFi HotStart ReadyMix (Roche Diagnostics, #KK2602) formulation which performs well for (A+T)-rich genomes like *P. falciparum*. Using mock samples of varying parasitemia, we searched for a reaction optimum in several full factorial experimental designs. For simplicity, we maintained a 2-step PCR program and 25 *µ*L reaction volume throughout, but explored varying 5 other factors (template DNA amount, number of cycles, primer concentration, polymerase concentration, and annealing temperature). By agarose gel, we observed that increasing the template amount increased sensitivity up to 8 *µ*L after which there was no clear benefit; and that a program with 35 cycles was more sensitive than 30 cycles (Supplementary Fig. 1a). A gradient PCR indicated an ideal annealing temperature of between 58–60 °C (Supplementary Fig. 1b). Increasing the amount of primer increased sensitivity but also increased visible background amplification, whereas increasing the amount of polymerase seemed to increase sensitivity without visibly affecting background (Supplementary Fig. 1c). Using this information, we selected four candidate optimal conditions, and sequenced seven mock samples under each (Supplementary Fig. 1d). From these, we selected the multiplex PCR conditions to maximize the percentage of reads mapping to *P. falciparum* malaria and mean coverage across amplicons. The final conditions were: 95 °C, 3 min; followed by 35-cycles of 98 °C 20 s, 60 °C 3 min; and a final at 60 °C for 10 min.

### NOMADS-MVP Sequencing Protocol

The complete laboratory protocol, including materials and primer sequences, is available online at protocols.io (English: NOMADS-MVP: Rapid Genomic Surveillance of Malaria; French: [Surveillance Génomique du Paludisme par la méthode Nanopore: Protocole Rapide NOMADS-MVP](https://www.protocols.io/view/surveillance-g-nomique-du-paludisme-par-la-m-thod-d64p9gvn)). In brief, 8 *µ*L of extracted DNA is used as template in a 25 *µ*L multiplex PCR using 15.5 *µ*L of Kapa HiFi HotStart ReadyMix (Roche Diagnostics, #KK2602) and 1.5 *µ*L of the NOMADS-MVP primer pool. Multiplex PCR products are cleaned using a 0.5X ratio of AMPure XP Beads (Beckman Coulter, A63881) and eluted in 15 *µ*L of nuclease-free water. DNA elute is quantified using the Qubit dsDNA HS Assay Kit (ThermoFisher Scientific, #Q33231) and between 200–800 ng of DNA is taken forward for barcoding and sequencing. We use Rapid Barcoding Kit 96 (SQK-RBK114.96) from Oxford Nanopore Technologies (ONT) following the associated ONT protocol with the following exceptions: we use 1 *µ*L (rather than 1.5 *µ*L) of each barcode during rapid barcoding; after barcoding we purify the pooled samples at a 0.5X ratio and elute in 15 *µ*L; we use 800 ng of DNA for adapter ligation, with a master mix of 1.0 *µ*L of RA and 2.3 *µ*L ABD (rather than 1.5 *µ*L RA, 3.5 *µ*L ADB). All sequencing was done using R10.4.1 flow cells on MinION Mk1B or Mk1D devices.

### Exclusion of selective whole genome amplification (sWGA)

We developed a modified version of the selective whole-genome amplification (sWGA) protocol^37^ that substituted the phi29 DNA polymerase (New England Biolabs, #M0269S) with EquiPhi29 (ThermoFisher Scientific, #A39390) as follows: 2 *µ*L EquiPhi29 10x buffer, 0.2 *µ*L 0.1M DTT, 2 *µ*L 500 *µ*M sWGA primer pool, 1 EquiPhi29 DNA polymerase, 12.8 *µ*L DNA; 45 °C 1hr, 60 °C 10min; dilute before PCR with 160 *µ*L nuclease-free water. We were able to obtain robust amplification with these conditions despite the substantial reduction in incubation time relative to phi29 DNA polymerase. However, despite both field and mock DBS samples typically yielding stronger bands on an agarose gel if sWGA was used, we saw no consistent improvement in the number of samples passing our quality control (≥8 amplicons with ≥ 50× mean coverage). In other words, samples that pass with using sWGA also passed without sWGA; and using sWGA to process samples did not rescue samples that failed without it. At the same time, we saw evidence that using sWGA skewed the proportions of strains within polyclonal infections, as has been documented previously^22^. Given that it also adds cost and complexity to the protocol, we removed it in August 2024.

### Real-time bioinformatics pipeline and dashboard

We developed a real-time bioinformatics pipeline and dashboard for nanopore sequencing, called *Nomadic*. The source code is publicly available on GitHub (https://github.com/JasonAHendry/nomadic) and user documentation is hosted on GitHub Pages (https://jasonahendry.github.io/nomadic/). *Nomadic* was designed for *P. falciparum* genomic surveillance with the NOMADS-MVP protocol, but was coded flexibily and supports other amplicon panels or organisms. Both the bioinformatics pipeline and dashboard are implemented in Python.

### Running a real-time analysis

Briefly, to run *Nomadic* (v0.5.0), a user first starts a nanopore sequencing run using *MinKNOW*, ensuring to: (1) assign a unique experiment name, hereafter referred to as `<expt_name>`; (2) enable real-time basecalling using either the High Accuracy (HAC) or Super Accurate (SUP) model; (3) enable sample demultiplexing with the appropriate barcoding kit (for NOMADS-MVP, SQK-RBK114.96). The user must prepare a comma-separated values (CSV) metadata table, which maps the barcodes to sample identifiers, and save it in a *Nomadic* workspace folder with the name `<expt_name>.csv.` The user can then launch *Nomadic* from a terminal window with the command `nomadic realtime <expt_name>`.

### Implementation and bioinformatics details

As sequencing proceeds, *MinKNOW* writes batches of basecalled reads to disk every ten minutes as FASTQ files. These are demultiplexed by *MinKNOW* such that each barcode possesses its own folder containing associated FASTQ files. *Nomadic* keeps track of the FASTQ files in these folders and, when a new FASTQ file has been generated for a given barcode, launches a per-sample bioinformatics pipeline to process it and update results for the sample. In brief, the pipeline maps reads to the *P. falciparum* 3D7 reference genome (PlasmoDB release 67) using *Minimap2*^52^ (v2.28), summarizes the output of mapping (i.e. fraction of reads mapped) and coverage of the target amplicons using *samtools* (v1.17), performs variant calling with *bcftools call*^38^ (v1.17) or *Delve* (v0.2.0), and annotates variants using *bcftools csq* (v1.17). Once a per-sample bioinformatics pipeline has been completed, a set of summary CSV files describing the results across all samples in the experiment is updated to reflect the incorporation of the new reads.

The dashboard runs in a separate thread and creates a graphical summary of these summary CSV files which updates every minute. The plots are interactive and users can hover over features of interest to open tooltips with additional information, zoom into areas of interest, or export static versions of the plots as PNG files. We implemented it using the Dash library (v2.17.1) from plotly. In our experience, sequencing and basecalling are often slower than the time *Nomadic* takes to process batches of FASTQ files, which allows the dashboard to remain up-to-date throughout the sequencing run. Once sequencing is completed, *MinKNOW* and *Nomadic* are stopped by the user. The final summary CSV files contain key results and are sufficient to reopen the dashboard, as described below.

### Viewing the dashboard for a previous experiment

The *Nomadic* dashboard can be reopened after a sequencing experiment is completed by running the command `nomadic dashboard <expt_name>` in a terminal window.

### Sequencing coverage summary statistics

We used summary files from *Nomadic* to compute per-sample summary statistics of sequencing coverage and evaluate performance across countries and parasitemia levels. In particular, we computed the mean coverage across all amplicons using the `mean_cov` column in the `summary.bedcov.csv` files produced by *Nomadic*. To summarise coverage uniformity across amplicons, we compute a fold-difference between the most- and least-abundant amplicon per sample, excluding the *hrp2/3* amplicons. In particular, defining *m*_*i* *j*_ as the mean coverage in sample *i* of amplicon *j*, and *a* is the set of all amplicons excluding *hrp2* and *hrp3*, then we compute:

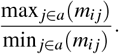

Excluding *hrp2* and *hrp3* makes the statistic robust to *hrp2/3* deletions; and in practice, for NOMADS-MVP *hrp2* or *hrp3* are rarely the lowest abundance amplicon in the absence of deletion and so do not contribute to the statistic’s value.

### Analysis of read lengths

As each experiment typically contains millions of reads, for computational simplicity we included only four experiments in this analysis: from Kenya (sequenced on 2024/07/30), Mali (2024/10/25), Ethiopia (2024/11/27) and a mock sample experiment (2024/11/26). For each experiment, we randomly sampled 20 barcodes (samples) without replacement from the those that: (i) had 8 or more amplicons with greater than 50× coverage; (ii) had >100 parasites/µL; (iii) did not use sWGA; (iv) were not positive or negative controls. For these barcodes, the BAM file generated by *Nomadic* was processed using *Pysam* (v0.22.1) to determine read lengths from the ‘query length’ field. Genomic regions of interest were delineated with a custom python script that converted gene codon numbers of interest into genomic coordinates using the *P. falciparum* 3D7 reference genome (PlasmoDB release 67) and associated general feature format (GFF) file, and then were manually verified using the Integrative Genome Viewer (IGV) (v2.5.0). We determined the percentage of reads overlapping by first using the intersect command from *bedtools*^53^ (v2.31) which enabled us to filter the BAM file to only reads overlapping the region of interest; and then comparing the number of mapped reads in this BAM file against the original BAM file using *samtools* (v1.17).

### Inference of *hrp2/3* deletions

We used a set of 219 previously published samples from Asella, Ethiopia to evaluate concordance between NOMADS-MVP and a gold-standard approach for *hrp2/3* deletion detection^42^. Recommended^43^ conventional PCRs for *hrp2* and *hrp3*^54^ were performed in duplicate and visualised by agarose gel electrophoresis. For discordant results, we called a sample positive if there was a clear band of the appropriate length in at least one of the replicates^43^. We excluded from analysis 46 samples that failed a *kelch13* PCR, suggesting low DNA quality or parasitemia and 24 samples that contained other *Plasmodium* species, leaving 149 samples. Of the 149 samples, 138 (92.6%) had 8 or more amplicons exceeding 50× coverage after NOMADS-MVP nanopore sequencing and were used for the comparison.

We predicted *hrp2/3* deletions from NOMADS-MVP sequencing data using a previously developed Bayesian statistical model^32^. In brief, it works as follows. For each sequencing run, the statistical model first estimates the rate of background contamination / barcode misclassification using included negative controls. Next, it estimates the quality of each sample and average variation in quality across samples using the mean coverage over all amplicons in the multiplex PCR, excluding the *hrp2* and *hrp3* amplicons. Finally, across all samples the posterior probability of *hrp2* and *hrp3* deletion are estimated separately using Markov Chain Monte Carlo (MCMC). Here, each chain was run for 50,000 iterations with a prior deletion probability of for *hrp2* and *hrp3*. Although a lower prior might be justified for hrp2 given prevalence of deletions in the region, in practice our estimates were insensitive to the prior. For Figure 5a, we directly show the posterior probabilities from the model. For Figure 5b, we are showing the *maximum a posteriori* (MAP) estimate of deletion. The statistical model and Bayesian MCMC were implemented in Python.

**Figure 5.**
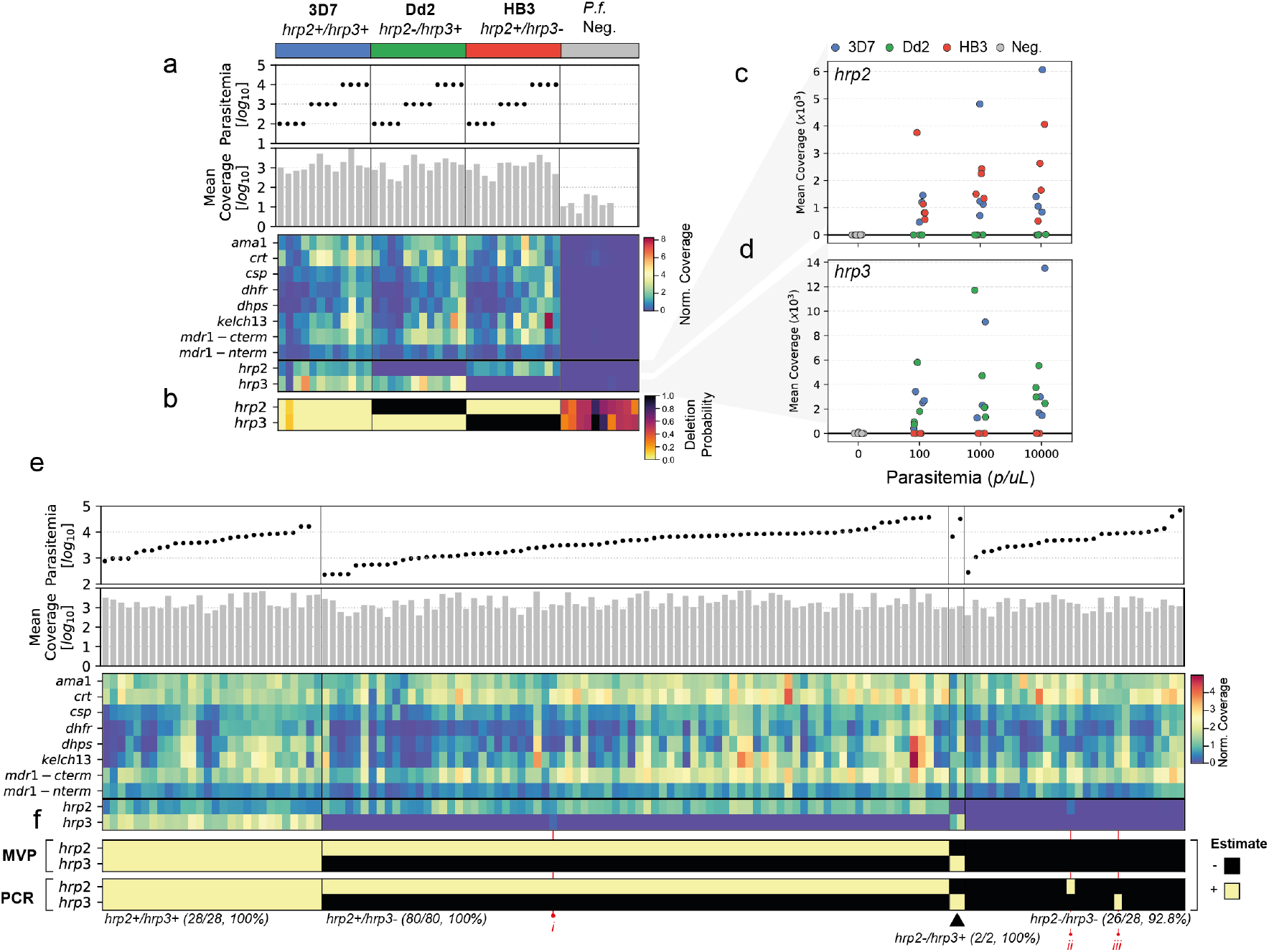
Validation of NOMADS-MVP *hrp2/3* deletion detection on mock and field DBS samples. (a) Data for 36 mock DBS samples of different *hrp2/3* genotypes and mock *P*.*f*. samples, which contain only human DNA (Methods). From top to bottom, subpanels show: lab strains used in mock samples; scatter plot of parasitemia; bar plot of mean coverage; heatmap with rows indicating different amplicons in NOMADS-MVP, columns indicating samples, and color indicating mean coverage, normalised to the mean coverage for the experiment. (b) Heatmap showing probability of deletion for *hrp2* and *hrp3* from statistical model. Note uncertainty (i.e. probability in 0.2–0.8) for negative controls, due to low or absence of coverage. (c) Scatter plot showing mean coverage over *hrp2* for all mock samples. (d) Same as (c), but for *hrp3*. (e) Same as (a), but showing data for 149 Ethiopian DBS samples. Vertical lines delineate *hrp2/3* deletion status as estimated by NOMADS-MVP. (f) Estimated *hrp2/3* deletion status by NOMADS-MVP (top subpanel) and the conventional PCR-based assays (bottom). Samples marked with numerals in red font indicate discrepancies (ii, iii) or residual coverage (i).

### Variant calling

We developed a novel variant caller, named *Delve*, to enable the detection of biallelic SNPs at low frequencies in polyclonal infections. *Delve* was implemented in Rust and is publicly available on GitHub (https://github.com/berndbohmeier/delve). Below we describe the statistical model, genotype inference, bias filtering, and model tuning. The notation used throughout this section is summarised in Table 2.

**Table 2.** Notations for *Delve*. Symbols representing model data and parameters are separated. Data is processed one site at a time; all symbols refer to a single genomic site. WSAF, within-sample alternative allele frequency; LRT, likelihood-ratio test.

| Data |  |  |
| --- | --- | --- |
| Symbol | Description |  |
| REF | Reference base | $\text{REF} \in \{A, T, C, G\}$ |
| ALT | Most common alternative base | $\text{ALT} \in \{A, T, C, G\} - \text{REF}$ |
| $D$ | Read depth | $D \in \mathbb{Z}^+$ |
| $j$ | Read index | $j \in \{1, 2, \dots, D\}$ |
| $b_j$ | Read base | $b_j \in \{\text{REF}, \text{ALT}\}$ |
| $q_j$ | Read base Phred-scaled quality score | $q_j \in \mathbb{Z}^+$ |
| $\epsilon_j$ | Read base error probability | $\epsilon_j \in [0, 1]$ |
| $v$ | Fraction of clones carrying ALT | $v \in [0, 1]$ |
| Parameters |  | Tuned Value |
| $D_{\min}$ | Minimum depth threshold after filtering | 20 |
| $Q_{\min}$ | Minimum base quality threshold | 20 |
| $v_0$ | Null-hypothesis for WSAF in LRT | 0.01 |
| $c_{LRT}$ | LRT threshold | 8 |
| $c_{ORT}$ | Strand-bias Odds Ratio Test threshold | 8 |

#### Statistical model

*Delve* assumes genomic sites are independent and calls SNPs at one site at a time. Reads overlapping a given site are indexed by *j* = 1, 2, …, *D*, where *D* is the sequencing depth at the site. Each read contributes a base, *b* _*j*_, which was aligned to the reference base REF ∈ {*A, T,C, G*} during read mapping, and an associated Phred-scaled base quality score, *q* _*j*_. *Delve* assumes sites are biallelic, and for each site retains only the reads carrying the reference base REF and most common alternative base, ALT ∈ {*A, T,C, G*} *−* REF; all other bases are treated as errors and discarded. Quality scores are taken from the basecaller, and then further adjusted using the Base Alignment Quality (BAQ) algorithm implemented in *samtools*^38^ to account for local alignment ambiguity. Afterwards, bases that have a quality score below the fixed threshold *Q*_min_ are discarded. If the depth after filtering is less than *D*_min_, no SNP call is made at the site.

A *P. falciparum* infection can consist of multiple clones at unknown proportions and, as a result, the fraction of clones carrying the alternative base, *ν*, can take on any value in the interval [0, 1]. We are interested in determining *ν* from the sequencing data. Assuming that reads are independent, the likelihood for *ν* is given by:

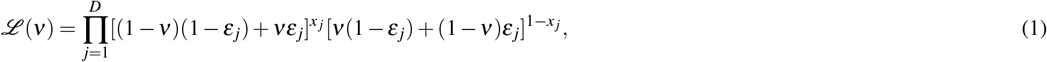

where *x* _*j*_ = 1 if *b* _*j*_ = REF, and *x* _*j*_ = 0 if *b* _*j*_ = ALT; and *ε* _*j*_ are the base error probabilities computed from the Phred-scaled quality scores, *ε* _*j*_ = *−*10 log_10_(*q*_*j*_).

This likelihood is a more general form of the genotype likelihood described by Li^55^ (implemented in *bcftools*) as well as by McKenna et al.^56^ (implemented in GATK). In those cases, the likelihood is parameterised in terms of a fixed genotype ploidy, represented by an integer. For example, in the diploid case, an organism can carry 0, 1, or 2 copies of the alternative allele, which, in the parameterisation above, corresponds exactly to *ν* = 0, *ν* = 0.5 or *ν* = 1.0. The key difference is that in this formulation we allow *ν* to vary continuously to reflect the unknown fraction of *P. falciparum* clones carrying the alternative base. For numerical stability, the likelihood is always evaluated in logarithmic space, yielding the log-likelihood *ℓ*(*ν*).

#### Genotype inference

The goal of inference is to learn both *ν*, the fraction of clones carrying the alternative base, and also use this *ν* to assign a genotype for the site. First, we determine the maximum likelihood estimate of 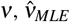, using Brent’s method bounded inside of the interval [0, 1]. We then use the likelihood ratio test (LRT) to evaluate the evidence for a SNP at the site. In particular, we test the null hypothesis that there is no variant, *H*_0_: *ν* ≤ *ν*_0_, versus the alternative hypothesis that there is a variant, *H*_1_: *ν > ν*_0_. In this way, to call a variant, we require that the 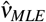 exceeds a small cut-off frequency *ν*_0_; in the absence of such a cut-off (i.e. if *ν*_0_ = 0), even the most subtle sequencing biases could eventually lead to a rejection of the null hypothesis, and spurious calling of a variant, as arbitrarily more reads were sequenced. Given these hypotheses, the LRT statistic is:

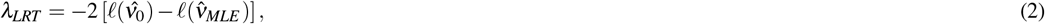

where 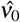 is the maximum likelihood estimate (MLE) for the alternative allele frequency *ν*, given the restriction *ν* ≤ *ν*_0_. We reject *H*_0_ if *λ*_*LRT*_ is greater than a threshold *c*_*LRT*_, which is set during model tuning. If the null hypothesis is rejected, we conduct a further likelihood ratio to distinguish between a homozygous alternative SNP, *H*_0_: *v* ≥ (1 *−ν*_0_), versus a heterozgyous alternative SNP, *H*_1_: *v <* (1 *−ν*_0_).

#### SNP filtering

*Delve* filters candidate SNPs if they exhibit excessive strand bias. In particular, we implemented the StrandOddsRatio test used in GATK^56^, with minor modifications. For each candidate SNP, we count the number of reference bases on the forward (*N*_*r*+_) and reverse strand (*N*_*r−*_), and the number of alternative bases on the forward (*N*_*a*+_) and reverse (*N*_*a−*_) strand. Using these counts, we calculate the odds ratio:

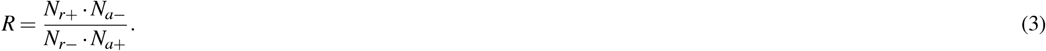

Following GATK, we make the ratio symmetric by calculating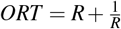.

A candidate SNP is filtered from the final SNP set if *ORT* exceeds a threshold *c*_*ORT*_. The rapid barcoding kit (SQK- RBK114.96) introduces strand coverage bias at amplicon edges, with the majority of coverage at the 5’ and 3’ ends of the amplicons deriving from the reverse and forward strands, respectively. This is an expected consequence of the transposome inserting barcodes uniformly across the amplicon and sequencing proceeding in a 3’ to 5’ direction. To accommodate this, we only apply the ORT filter to SNPs that have a heterozygous alternative genotype call; in this way avoiding filtering homozygous alternative SNPs with strand coverage bias due to the barcoding chemistry.

#### Model tuning

In total, *Delve* is tuned by five parameters (Table 2). We set the base-quality filter *Q*_min_ by examining the distribution of quality scores after applying the BAQ algorithm, observing a long tail of lower quality bases beginning at *Q* = 20 and comprising approximately 14% of all bases. We explored varying *v*_0_, *c*_*LRT*_, *c*_*ORT*_ to maximise recall, under the constraint of maintaining near-perfect precision in the 1,000 and 10,000 parasites/µL mock DBS samples. The minimum depth filter *D*_min_ was disabled during the downsampling analysis. Otherwise, we set *D*_min_ = 20; although in principle, the *c*_*LRT*_ threshold already implicitly sets a threshold on the amount of sequencing depth (and base qualities) required to make a variant call.

### SNP calling accuracy evaluation

#### Creating a set of true variants

3D7, Dd2, HB3 have been sequenced using Pacific Bioscience Sequencing SMRT technology and high-quality FASTA sequences available on *PlasmoDB*^57^. To identify variants in these assemblies with respect to the 3D7 reference genome, we simulated high-quality (Phred 60) error-free reads *in silico* from the FASTA files, mapped them to the 3D7 reference genome with *minimap2* (v2.28), and then identified variants using the *bcftools*^38^ (v1.22) `mpileup` and `call` commands. We simulated 60 error-free reads, half forward and half reverse strand, for each amplicon in our NOMADS-MVP panel by extracting the FASTA sequence spanning +/-4 kbp of the target, based on GFF files for 3D7, Dd2 and HB3. This procedure resulted in truth VCF files for each of the clonal strains. We created truth VCF files for the two-strain mixtures by combining allelic depth information from the clonal strain VCF files at the correct proportions given the laboratory mixture, and updating the genotype call accordingly.

For each Cambodian strain, information about the *Kelch13* amino acid changes were available from BEI resources (IPC3445, C580Y; IPC4912, I543T; IPC5202, R539T); however, we found no information about corresponding nucleotide changes. Therefore, we performed Sanger sequencing using the NOMADS-MVP *kelch13* amplicon as a singleplex PCR to determine the *kelch13* nucleotide sequence for each strain. We used *tracy*^58^ to convert the chromatogram file into a VCF with the `tracy decompose` command.

#### Evaluating accuracy

For all SNP calling accuracy evaluations we used the Super Accurate (SUP) basecalling model in *MinKNOW* (v24.06.16) to generate FASTQ files, and then ran *Nomadic* in real-time to map reads and produce BAM files. We called variants using both the *bcftools* (v1.22) `call` command and *Delve* (v0.5.0). We used *hap*.*py*^59^ from a *Docker* image (jmcdani20/hap.py:v0.3.12) to compute measures of variant calling accuracy in comparison to the truth VCF files described above. In particular, we used three measures of SNP calling performance: (i) the precision, *TP/*(*TP* + *FP*), where *TP* is the number of true positive SNP calls (homozygous or heterozygous alternative), and *FP* is the number of false-positive SNP calls; (ii) the recall of true homozygous alternative SNPs, *T P*_*homo*_*/*(*T P*_*homo*_ + *FN*_*homo*_), where *T P*_*homo*_ and *FN*_*homo*_ are the number of true homozygous alternative SNPs called and missed, respectively; (iii) the recall of true heterozgyous alternative SNPs, *T P*_*het*_*/*(*T P*_*het*_ + *FN*_*het*_), which is defined analogously with (ii). We excluded from the analysis: (i) low-complexity regions of the Pf3D7 reference genome, identified using *sdust* with default parameters; (ii) the central repeat region of *csp*; (iii) the *hrp2* and *hrp3* amplicons, as their primary utility is for deletion identification and they consist largely of low-complexity repetitive sequence. This left a total of 6,681 bp for evaluation in each mock DBS sample. For amplicons where no variants existed for a particular mock sample (e.g. *Kelch13* for 3D7, Dd2 and HB3), the precision and recall are considered to be undefined; unless false-positive mutations are present, in which case the precision is zero. The within-sample alternative allele frequencies (WSAFs) plotted in Figure 4a,c were calculated directly from the allelic depths.

#### Downsampling analysis

For the downsampling analysis, we included all mock DBS samples that: (i) were at 10,000 or 1,000 parasites/µL; (ii) had greater than 500× coverage for all amplicons. This set included a total of 72 mock DBS samples (33 with 1,000 parasites/µL and 39 with 10,000 parasites/µL). We excluded the mock DBS samples with 100 parasites/µL because the majority (62.2%, 28/45) did not have greater than 500× coverage for all amplicons. For each mock DBS sample, we randomly downsampled the sequencing reads to a specific mean coverage for each amplicon, using *samtools*^55^ (v1.22). To achieve this, we split each mock sample’s BAM file into 10 per-amplicon BAM files, each containing reads mapping to one of the NOMADS-MVP targets. We then downsampled these per-amplicon BAM files independently, with the `samtools view` command and `-s/--subsample` flag, to achieve a target mean coverage level, before concatenating them back into a single, per-sample BAM file; now with all amplicons having the target coverage. We downsampled to a mean coverage of 500, 400, 300, 200, 150, 100, 75, 50, and 25×, in triplicate for each mock DBS sample and coverage level, producing a total of 1,944 BAM files for variant calling with *Delve* (72 samples by 9 coverage levels, in triplicate).

## Acknowledgements

The NOMADS project is funded by the Bill Melinda Gates Foundation (INV-048316 to M.M., D.J.B. and J.A.H.). Sample collection and *hrp2/3* deletion investigation in Ethiopia was supported by the ARSUNA project, funded by the German Federal Ministry for Economic Cooperation and Development and the Else Kröner-Fresenius Foundation (EKFS) via the Hospital Partnerships Programme (project 21Ac01060). Work in Côte d’Ivoire was funded by a grant from the Global Health Protection Program (Federal Ministry of Health of Germany, grant number ZMII2-2523GHP029) and the WHO Hub for Pandemic Preparedness. We thank Bob Verity for helping to estimate the number of samples required to detect mutations spreading across Africa with adequate power; Olaf Kostbahn, Wendy Vienneau, and Sossena Assefa for critical financial and legal administration; and Estée Török for fruitful scientific discussions, feedback and encouragement throughout the project’s duration. We are grateful to all those who participated in the studies conducted across Africa, especially the health workers and patients who made the project possible.

## Author contributions statement

M.M.: Conceptualization, Methodology, Validation, Investigation, Data curation, Funding acquisition. K.M.: Methodology, Validation, Investigation. B.B.: Methodology, Software, Validation, Formal analysis. M.C.: Investigation. W.V.L.: Investigation, Resources, Data curation. B.M.: Investigation. A.G.: Investigation. A.O.: Investigation. S.S.: Investigation. N.J.Y.: Investigation. D.S.: Investigation. B.N.: Data curation. D.Z.: Investigation. Y.N.G.T.: Investigation. F.O.: Investigation, Data curation. A.Z.: Investigation. E.A.A.: Investigation. C.C.: Investigation. S.O.: Investigation. B.O.: Investigation. M.N.: Resources. D.K.G.: Resources. T.F.: Resources. T.B.T.: Resources. R.O.: Investigation. E.S.: Investigation. Y.S.: Resources. C.N.: Resources. O. Ouedraogo.: Resources. K.O.: Resources. O.Opaleye.: Resources. A.Olowe.: Resources. M.G.: Project administration. C.D.: Supervision. G.S.: Resources, Data curation, Supervision. F.P.M.: Resources, Supervision. S.P.: Resources, Supervision. A.D.: Resources, Supervision. I.S.: Resources, Supervision. D.N.: Resources, Supervision. O.Ojurongbe.: Resources, Supervision. S.K.: Resources, Supervision. Y.P.S.: Conceptualization, Resources, Data curation, Supervision. J.S.S.: Resources, Data curation, Supervision. M.H.: Resources, Supervision. D.J.B.: Conceptualization, Data curation, Supervision, Funding acquisition. J.A.H.: Conceptualization, Methodology, Software, Validation, Formal analysis, Data curation, Supervision, Funding acquisition, Writing - Original Draft. All authors reviewed and approved the final version of the manuscript.

## Additional information

The authors declare no competing interests. **CDC Disclaimer:** The opinions expressed by the authors do not necessarily reflect the official views of the CDC or the U.S. Department of Health and Human Services.

## Notes

### Competing Interest Statement

The authors have declared no competing interest.

### Summary of Updates

Updated to include analysis of SNP calling in polyclonal infections; description of new variant caller in Methods; two new results sections; new Figure 4; new Supplementary Table 4; removed previous Figure 4 and Supplementary Figure 6 as these are superseded by the new analysis.

