## Supplementary Materials for "Continental-scale genomic surveillance of *Plasmodium falciparum* malaria with rapid nanopore sequencing"

| Item | Manufacturer | Catalog # | Pack Size | Unit Cost ($) | Units | Cost ($) |
|---|---|---|---|---|---|---|
| 1.5 ml DNA LoBind tubes | Eppendorf | 0030 108.051 | 250 tubes | 18.97 | 1 | 18.97 |
| 100ml reservoir | ThermoFisher | 10125707 | 100 case | 248 | 1 | 248.00 |
| Adhesive seal sheets | Greiner | 676001 | 100 seals | 94 | 2 | 188.00 |
| Gloves | Handsafe | GN90-L | 2000 gloves | 88.01 | 3 | 264.03 |
| PCR strip tubes | ThermoFisher | AB2005 | 2000 wells | 354 | 1 | 354.00 |
| Plates 96 well (skirted) | ThermoFisher | AB-0800 | 25 plates | 115.94 | 7 | 811.58 |
| Qubit Assay Tubes | Invitrogen | Q32856 | 500 tubes | 111.41 | 5 | 557.05 |
| Sharpie markers | Stabilo | 01-19-2763 | 10 per pack | 15.89 | 1 | 15.89 |
| Tips 10ul | – | – | 4800 tips | 396.29 | 1 | 396.29 |
| Tips 200ul | – | – | 4800 tips | 396.29 | 1 | 396.29 |
| Tips 200ul (Multichannel) | – | – | 960 tips | 192.32 | 13 | 2500.16 |
| Tips 20ul | – | – | 4800 tips | 396.29 | 1 | 396.29 |
| Tips 20ul (Multichannel) | – | – | 960 tips | 192.32 | 26 | 5000.32 |
| Tips 1000ul | – | – | 3072 tips | 358.54 | 1 | 358.54 |
| **Total (plasticware)** | | | | | | **10,948.32** |
| **Cost per sample (plasticware)** | | | | | | **5.47** (26.2%) |
| Agarose | ThermoFisher | 16500500 | 500 gram | 675 | 1 | 675.00 |
| AMPure XP beads | Beckman Coulter | A63881 | 60 ml | 1485.20 | 1 | 1485.20 |
| DNA ladder (1kb) | ThermoFisher | SM0314 | 100 lanes | 64.99 | 2 | 129.98 |
| Ethanol | Sigma | E7024-500ML | 500 ml | 42.61 | 2 | 85.22 |
| Flow Cell Priming Kit | ONT | EXP-FLP004 | 6 rxns | 35.04 | 4 | 140.16 |
| Flow Cell Wash Kit | ONT | EXP-WSH004 | 6 washes | 99.10 | 4 | 396.40 |
| Gel loading dye | NEB | B7024S | 4 ml | 58.86 | 1 | 58.86 |
| KAPA HIFI ReadyMix | Roche | KK2602 | 6.25 ml | 1064.21 | 6 | 6385.26 |
| MinION Flow Cell (R10.4.1) 24 pack | ONT | FLO-MIN114 | 24 flow cell | 14400 | 1 | 14400.00 |
| Nuclease free water | Invitrogen | AM9937 | 500 ml | 155.19 | 1 | 155.19 |
| PCR Primers: MVP | IDT | N/A | 3000 rxn | 120 | 1 | 120.00 |
| Qubit 1x dsDNA HS Assay Kit | ThermoFisher | Q33231 | 500 rxn | 444.32 | 5 | 2221.60 |
| Rapid Barcoding Sequencing Kit 96 | ONT | SQK-RBK114.96 | 12 rxn | 990.99 | 4 | 3963.96 |
| TE buffer (1X) pH 8.0 low EDTA | ThermoFisher | J75793.AP | 500 ml | 120 | 1 | 120.00 |
| **Total (reagents)** | | | | | | **30,893.88** |
| **Cost per sample (reagents)** | | | | | | **15.45** (78.3%) |
| **Total** | | | | | | **41,842.00** |
| **Cost per sample** | | | | | | **20.92** (100.0%) |

**Supplementary Table 1: Itemized list of plasticware and reagents required for NOMADS-MVP.** 'Units' were calculated to allow for processing 2000 samples. A 10% mastermix overage was included where relevant. We assumed 48 samples/run and 2 runs per flow cell. All costs are in USD and were taken from quotes from Carramore International Limited (`https://www.carramore.com/`) for shipments to Africa made in 2025. No specific manufacturer for pipette tips is recommended, as they should be selected for compatibility with available pipettes. The cost of DNA extraction is not included.

| Application | Item | Manufacturer | Catalogue # | Item Cost ($) |
|---|---|---|---|---|
| DNA quantification | Qubit 4 Fluorometer | ThermoFisher | Q33238 | $4,400 |
| Sequencing | MinION | ONT | Mk1D† | $3,000 |
| Process data | Laptop with 1 Tb SSD, Nvidia GPU (16Gb RAM), i7 or equivalent processor, 16 Gb RAM | - | - | $3,000 |
| Store data | 5 Tb Storage Drive | - | - | $200 |
| DNA clean-up | Plate Magnet | ThermoFisher | 12027 | $1,100 |
| DNA clean-up | Tube Magnet | ThermoFisher | 12321D | $800 |
| **Total** | | | | **$12,500** |

**Supplementary Table 2: Itemized list of equipment required for NOMADS-MVP.** Main equipment for NOMADS-MVP is shown, basic laboratory equipment (PCR machine, vortex, centrifuge) is excluded. †The Mk1D will soon replace the Mk1B; note this cost includes a package of starting reagents. All costs are in USD and were taken from Carramore International Limited (`https://www.carramore.com/`). No specific recommendations are made for the laptop or external hard drive.

| Protocol Name | Platform | Incubation Time (mins) | Pipette Steps | Per-sample cost (USD) | Reference |
|---|---|---|---|---|---|
| MAD$^4$HatTeR | Illumina | 205 | 404 | $12–25 | *Aranda-Díaz et al.* [1] |
| *Pf*-SMARRT | Illumina | 340 | 565 | - | *Sadler et al.* [2] |
| NOMADS8/16 | ONT | 1400 | 292 | $25 | *de Cesare et al.* [3] |
| NOMADS-MVP | ONT | 206 | 172 | $20.92 | This study |

**Supplementary Table 3: Comparison of costs and complexity of NGS protocols for *P. falciparum* genomic surveillance.** 'Incubation Time' gives total incubation time across the entire protocol, not including hands-on pipetting time. For pipetting steps, we counted the number of steps from extracted DNA to sequencing for a batch of 48 samples, assuming access to an 8-channel pipette and that master mixes are pre-aliquoted into strip-tubes to minimize 96-well plate pipetting. Qubit-related steps were excluded as they are often optional and/or not done for all samples for convenience. One pipetting step includes both aspirating and dispensing a reagent or sample. Breakdown of per-sample costs for this study is available in STable 1. For other studies, we used per-sample costs from the indicated reference where available.

**Accuracy, % (TP/TN/FP/FN)**

| Samples with ≥1,000 parasites/$\mu$L | | | | | |
|---|---|---|---|---|---|
| Marker | Clonal | 20% | 10% | 5% | 2.5% |
| *crt* K76T | 100.0 (6/18/0/0) | 100.0 (12/6/0/0) | 100.0 (12/6/0/0) | 100.0 (12/6/0/0) | 72.2 (7/6/0/5) |
| *dhfr* N51I | 100.0 (6/18/0/0) | 100.0 (12/6/0/0) | 100.0 (12/6/0/0) | 100.0 (12/6/0/0) | 72.2 (7/6/0/5) |
| *dhfr* C59R | 100.0 (6/18/0/0) | 100.0 (12/6/0/0) | 100.0 (12/6/0/0) | 100.0 (12/6/0/0) | 55.6 (4/6/0/8) |
| *dhfr* S108N | 100.0 (12/12/0/0) | 100.0 (18/0/0/0) | 100.0 (18/0/0/0) | 100.0 (18/0/0/0) | 94.4 (17/0/0/1) |
| *dhps* S436F | 100.0 (6/18/0/0) | 100.0 (12/6/0/0) | 100.0 (12/6/0/0) | 100.0 (12/6/0/0) | 94.4 (11/6/0/1) |
| *dhps* A437G | 100.0 (12/12/0/0) | 100.0 (18/0/0/0) | 100.0 (18/0/0/0) | 100.0 (18/0/0/0) | 88.9 (16/0/0/2) |
| *dhps* A613S | 100.0 (6/18/0/0) | 100.0 (12/6/0/0) | 100.0 (12/6/0/0) | 100.0 (12/6/0/0) | 77.8 (8/6/0/4) |
| *kelch13* R539T | 100.0 (2/22/0/0) | - | - | - | - |
| *kelch13* I543T | 100.0 (2/22/0/0) | - | - | - | - |
| *kelch13* C580Y | 100.0 (2/22/0/0) | - | - | - | - |
| *mdr1* N86Y/F | 100.0 (6/18/0/0) | 100.0 (12/6/0/0) | 94.4 (11/6/0/1) | 77.8 (8/6/0/4) | 44.4 (2/6/0/10) |
| *mdr1* Y184F | 100.0 (6/18/0/0) | 100.0 (12/6/0/0) | 100.0 (12/6/0/0) | 100.0 (12/6/0/0) | 88.9 (10/6/0/2) |
| *mdr1* N1042D | 100.0 (6/18/0/0) | 100.0 (12/6/0/0) | 100.0 (12/6/0/0) | 100.0 (12/6/0/0) | 88.9 (10/6/0/2) |
| Samples with 100 parasites/$\mu$L | | | | | |
| Marker | Clonal | 20% | 10% | 5% | 2.5% |
| *crt* K76T | 100.0 (3/6/0/0) | 100.0 (6/3/0/0) | 100.0 (6/3/0/0) | 77.8 (4/3/0/2) | 66.7 (3/3/0/3) |
| *dhfr* N51I | 100.0 (3/6/0/0) | 100.0 (6/3/0/0) | 100.0 (6/3/0/0) | 55.6 (2/3/0/4) | 55.6 (2/3/0/4) |
| *dhfr* C59R | 100.0 (3/6/0/0) | 100.0 (6/3/0/0) | 100.0 (6/3/0/0) | 55.6 (2/3/0/4) | 55.6 (2/3/0/4) |
| *dhfr* S108N | 100.0 (6/3/0/0) | 100.0 (9/0/0/0) | 100.0 (9/0/0/0) | 66.7 (6/0/0/3) | 77.8 (7/0/0/2) |
| *dhps* S436F | 100.0 (3/6/0/0) | 100.0 (6/3/0/0) | 88.9 (5/3/0/1) | 44.4 (1/3/0/5) | 55.6 (2/3/0/4) |
| *dhps* A437G | 100.0 (6/3/0/0) | 100.0 (9/0/0/0) | 88.9 (8/0/0/1) | 55.6 (5/0/0/4) | 55.6 (5/0/0/4) |
| *dhps* A613S | 100.0 (3/6/0/0) | 100.0 (6/3/0/0) | 88.9 (5/3/0/1) | 33.3 (0/3/0/6) | 55.6 (2/3/0/4) |
| *mdr1* N86Y/F | 100.0 (3/6/0/0) | 88.9 (5/3/0/1) | 55.6 (2/3/0/4) | 66.7 (3/3/0/3) | 33.3 (0/3/0/6) |
| *mdr1* Y184F | 100.0 (3/6/0/0) | 100.0 (6/3/0/0) | 100.0 (6/3/0/0) | 77.8 (4/3/0/2) | 66.7 (3/3/0/3) |
| *mdr1* N1042D | 100.0 (3/6/0/0) | 100.0 (6/3/0/0) | 100.0 (6/3/0/0) | 88.9 (5/3/0/1) | 88.9 (5/3/0/1) |

**Supplementary Table 4: Evaluating the accuracy of antimalarial resistance marker identification across mock samples.** Rows display antimalarial resistance markers and columns group samples by minor clone frequency. Each cell shows the accuracy with which the marker was detected across samples and in the parentheses the count of true-positives (TP), true-negatives (TN), false-positives (FP), and false-negatives (FN). Accuracy is defined as $(TP + TN)/(TP + TN + FP + FN)$. Mock samples with 10,000 parasites/$\mu$L and 1,000 parasites/$\mu$L are grouped at top; and mock samples with 100 parasites/$\mu$L are shown at bottom. Only markers were at least one mock sample carries the mutation are shown. Note that the Cambodian strains carrying the *kelch13* markers were only prepared as clonal mock samples, due to the difficulties of making polyclonal samples accurately from DNA.

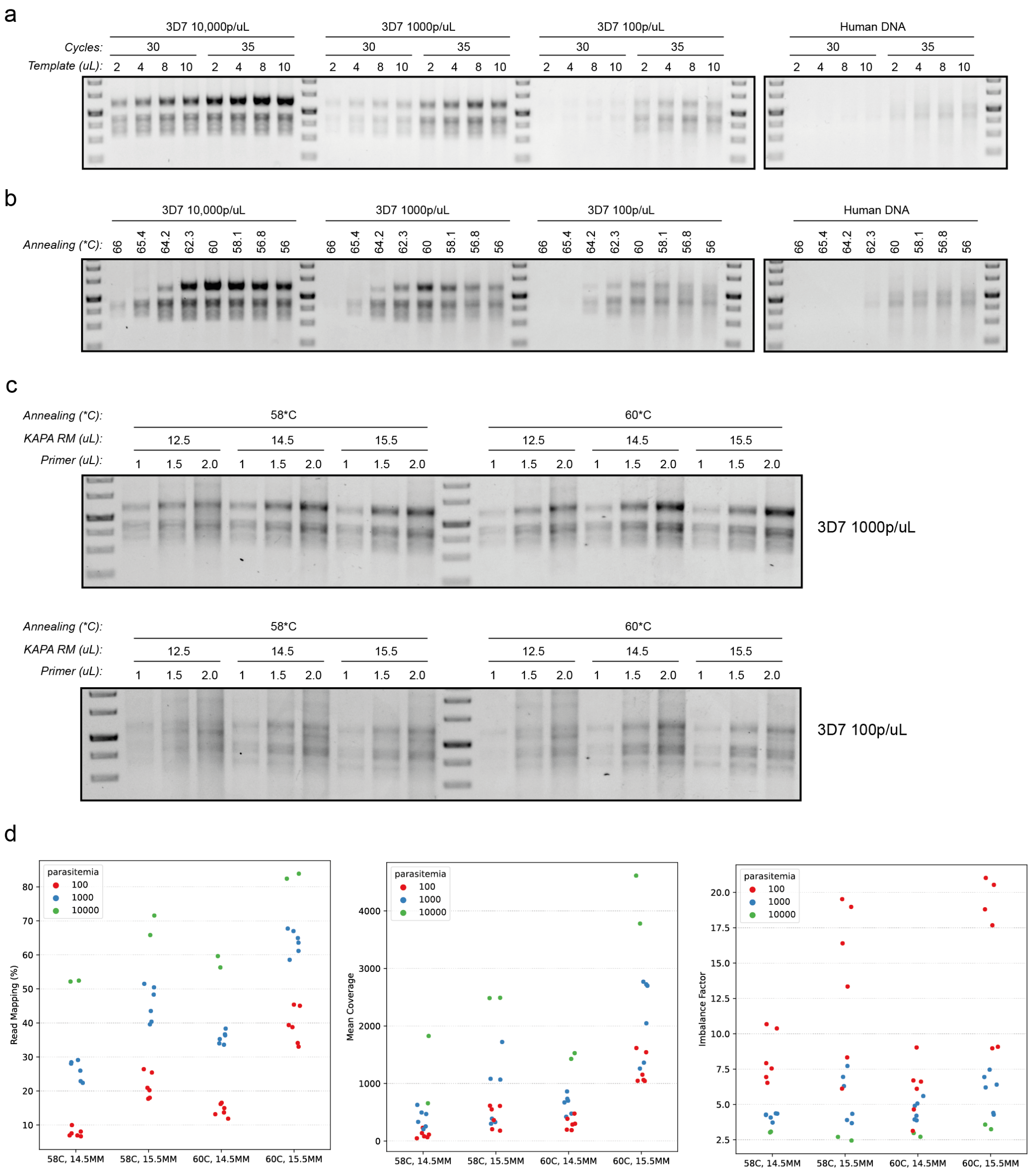


**Supplementary Figure 1: Optimising multiplex PCR conditions for NOMADS-MVP**. We optimised the conditions of the NOMADS-MVP multiplex PCR using mock samples. (a) Varying number of PCR cycles and template DNA amount. Total reaction volume was fixed at 25 $\mu$L. Based on these results, we used 8 $\mu$L template DNA and 35-cycles for experiments in panels (b) and (c). (b) Temperature gradient for the annealing/extension step. We restricted to a two-step PCR program with a denaturation step (98°C, 30 seconds) and a combined annealing and extension step (56–66°C temperature gradient, 3 mins). (c) Full factorial experiment varying annealing temperature (58°C, 60°C), KAPA HiFi ReadyMix volume (12.5, 14.5, 15.5 $\mu$L) and primer amount (1, 1.5, 2.5 $\mu$L) for two different mock samples (3D7 at 1000 parasites/$\mu$L and 100 parasites/$\mu$L). The KAPA HiFi ReadyMix is a 2X formulation so 12.5 $\mu$L is the manufacturer recommended concentration for a 25 $\mu$L reaction. Here, we saw boosted sensitivity at 15.5 $\mu$L without increased background. (d) Sequencing summary statistics for most promising conditions from (c) across several mock samples at different parasitemia.

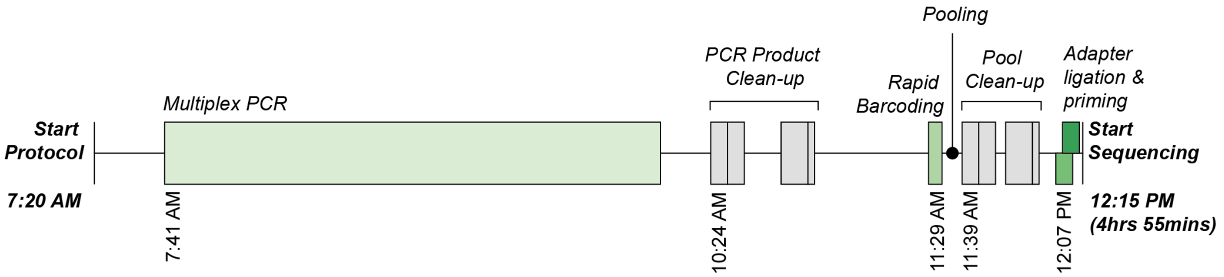


**Supplementary Figure 2: Diagram of NOMADS-MVP laboratory protocol timeline.** The diagram shows protocol time for a batch of 48 mock DBS samples sequenced at the Max Planck Institute for Infection Biology. Each rectangular box delineates an incubation step, and the lines connecting them were occupied with hands-on laboratory manipulation. Major steps are annotated. Each DNA clean-up step (grey) has four incubations: DNA binding to the AMPure XP magnetic beads; beads binding to the magnet; DNA elution from magnetic beads; removing beads from elute. We generated the timeline by keeping record of the start time of the experiment, the start time of each incubation step, and the start time for sequencing. The majority of the hands-on time occurs after PCR product clean-up and before rapid barcoding and is associated with running an agarose gel and quantifying DNA concentration using the Qubit Flourometer.

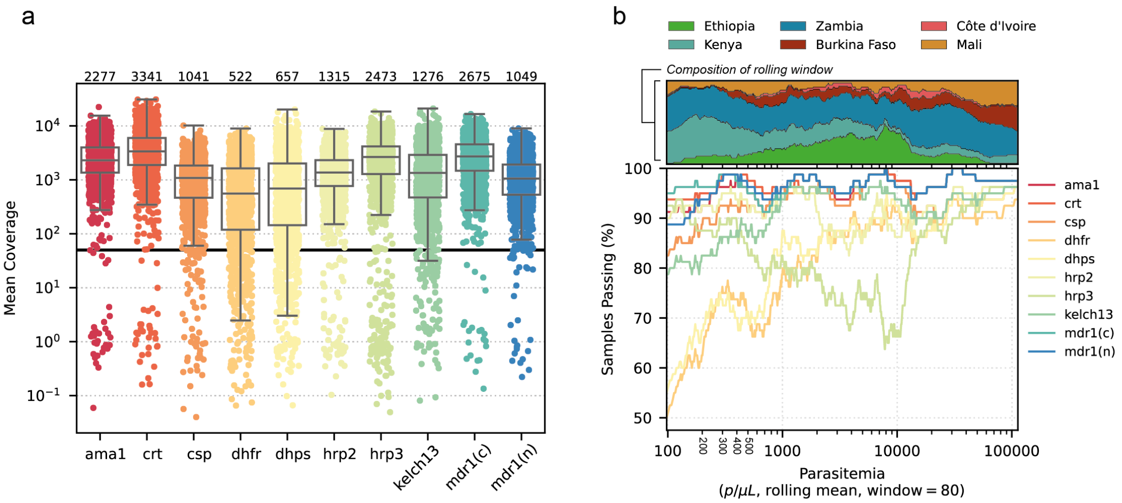


**Supplementary Figure 3: Performance of individual NOMADS-MVP amplicons.** (a) Boxplots showing mean coverage by amplicon across a total of 1283 DBS samples (1154 field, 129 mock) processed with NOMADS-MVP. The median abundance for each amplicon across all samples is annotated above. *crt* and *ama1* have the highest abundance and are the shortest amplicons. (b) Relationship between parasitemia and samples passing (%) grouped by amplicon. Field samples were ordered from lowest to highest parasitemia, and with a window size of 80 samples a rolling mean of the parasitemia (parasites/$\mu$L) was computed (x-axis) and the percentage of samples with $\geq 50\times$ coverage for each amplicon (y-axis) was computed to construct the lines. At lower parasitemias the pass rate of *dhfr* and *dhps* tends to decline. The dip in *hrp3* around 10,000 parasites/$\mu$L is due to a large proportion of samples from Ethiopia carrying *hrp3* deletions. The top pane displays what proportion (y-axis) of the 80 samples were from each country (by color) across the rolling windows (x-axis). Only field samples were included.

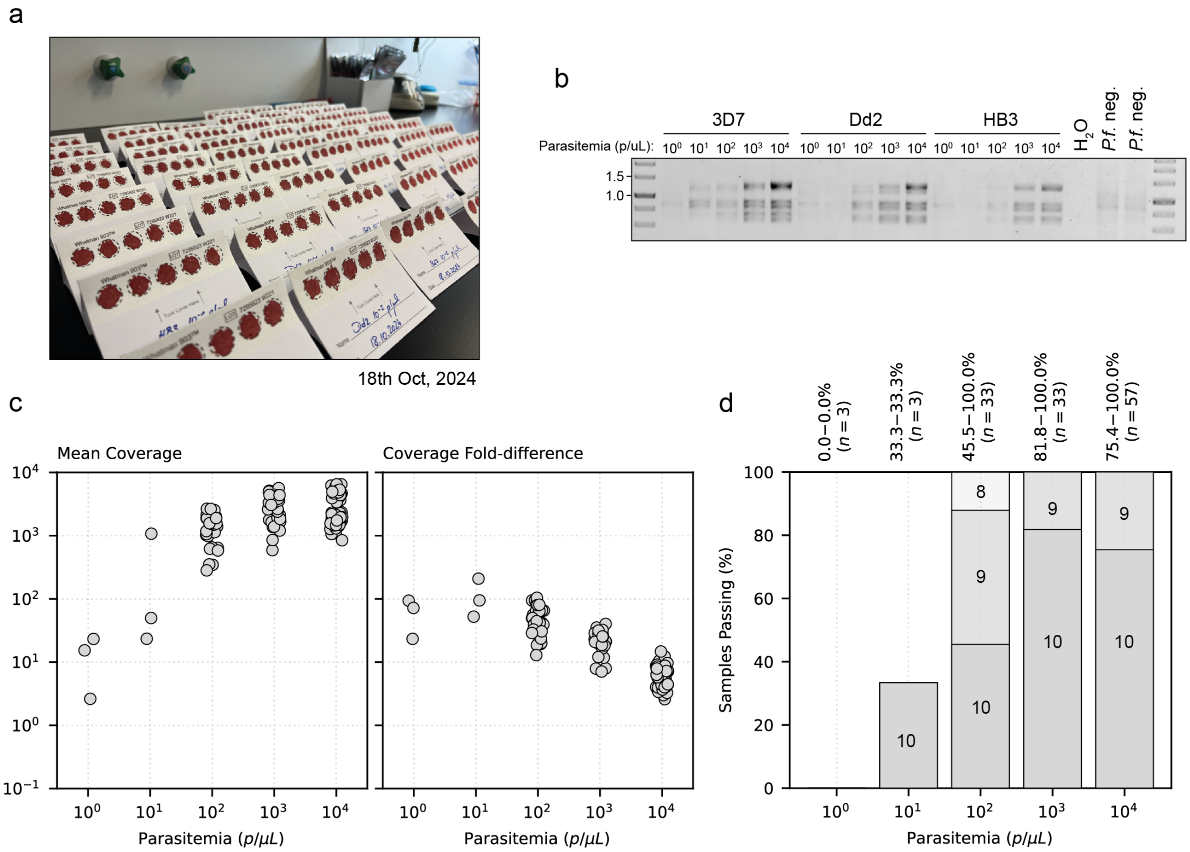


**Supplementary Figure 4: Performance of NOMADS-MVP on mock samples**. Mock DBS samples were created by culturing laboratory strains of *P. falciparum* and mixing them with human whole blood at different parasitemias. (a) Photo of the mock DBS generated for this manuscript. Each filter paper contains 5 spots of the same mock sample. (b) Agarose gel image of NOMADS-MVP performance on clonal mock DBS samples from 3D7, Dd2 and HB3 ranging from 1 to 10,000 parasites/$\mu$L. Negative controls of only water and human DNA are also included. (c) Scatterplots of mean coverage across and coverage fold-difference (Methods) for all mock DBS samples sequenced. (d) Barplot of percentage of mock samples passing across parasitemia. Note that 100% of mock DBS at 100 parasites/$\mu$L had at least 8 amplicons with $\geq 50\times$ coverage.

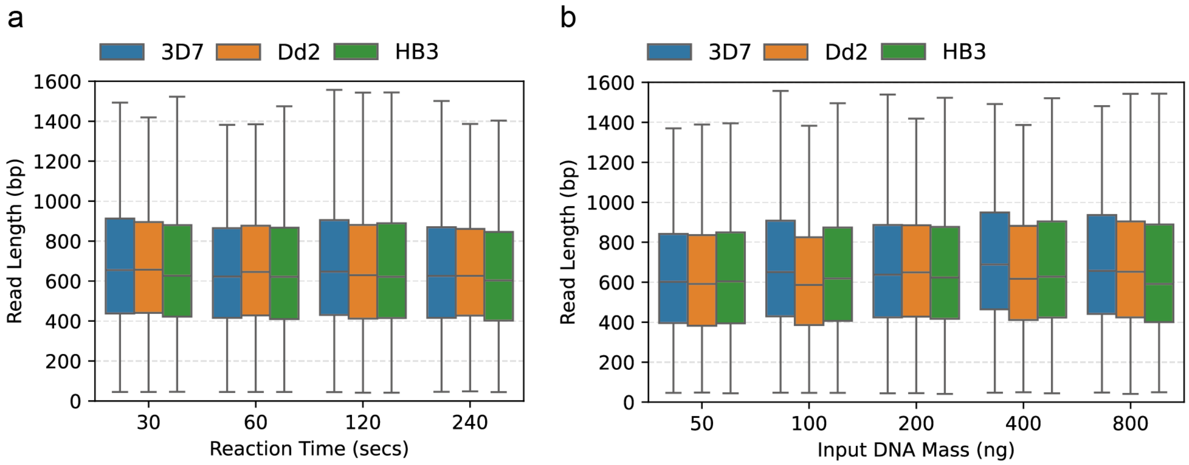


**Supplementary Figure 5: Evaluating the influence of the barcoding reaction on read lengths**. Rapid Barcoding Kit (SQK-RBK114.96) from ONT fragments reads during barcoding and we evaluated whether modulating either the (a) reaction time or (b) input DNA mass influenced read lengths (y-axis). To isolate the effect of the tagmentation reaction, multiple NOMADS-MVP multiplex PCRs were pooled for 3D7, Dd2 and HB3 at 10,000 parasites/$\mu$L. Aliquots of these pools were then used to conduct rapid barcoding for different reaction times or input DNA mass (ng) amounts. The recommended reaction from ONT uses 200ng of input DNA and a 120 second incubation. We observe little effect on read length when these are varied.

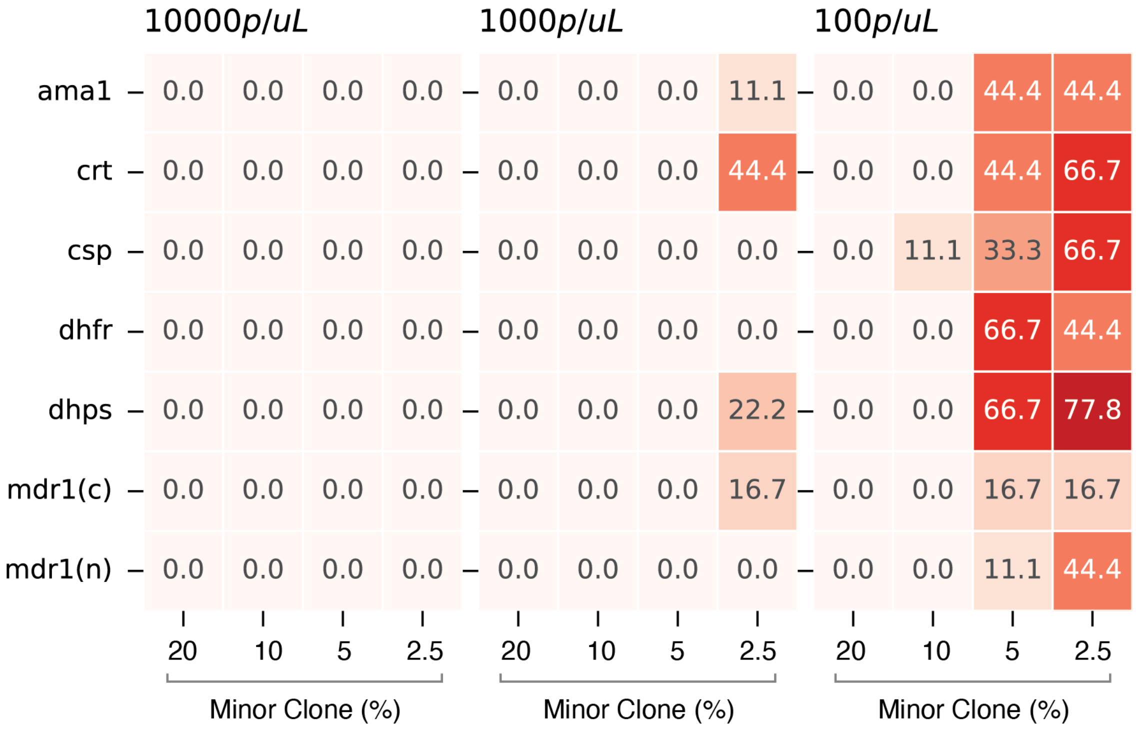


**Supplementary Figure 6: Assessing the loss of haplotype signal at low parasitemia levels and minor clone proportions**. Heatmap showing the percentage (%) of samples in which the minor clone haplotype generated no within-sample alternative allele (WSAF) signal in the sequencing reads. The data were grouped by parasitemia level (subpanels), amplicon (y-axis), and minor clone proportion (x-axis). For each amplicon and laboratory stain mixture, we determined the set of expected (true) heterozygous SNPs for the minor clone; if greater than 90% of these SNPs had a WSAF less than 0.5% or greater than 99.5%, we classified the haplotype as generating no signal. Notice that at 100 parasites/$\mu$L, a considerable fraction of samples generate no signal for the minor clone haplotype across all amplicons. Only amplicons with true heterozygous SNPs were included in the analysis.
